# Reference transcriptomes of porcine peripheral immune cells created through bulk and single-cell RNA sequencing

**DOI:** 10.1101/2021.04.02.438107

**Authors:** Juber Herrera-Uribe, Jayne E. Wiarda, Sathesh K. Sivasankaran, Lance Daharsh, Haibo Liu, Kristen A. Byrne, Timothy P.L. Smith, Joan K. Lunney, Crystal L. Loving, Christopher K. Tuggle

## Abstract

Pigs are a valuable human biomedical model and an important protein source supporting global food security. The transcriptomes of peripheral blood immune cells in pigs were defined at the bulk cell-type and single cell levels. First, eight cell types were isolated in bulk from peripheral blood mononuclear cells (PBMCs) by cell sorting, representing Myeloid, NK cells and specific populations of T and B cells. Transcriptomes for each bulk population of cells were generated by RNA-seq with 10,974 expressed genes detected. Pairwise comparisons between cell types revealed specific expression, while enrichment analysis identified 1,885 to 3,591 significantly enriched genes across all 8 cell types. Gene Ontology analysis for the top 25% of significantly enriched genes (SEG) showed high enrichment of biological processes related to the nature of each cell type. Comparison of gene expression indicated highly significant correlations between pig cells and corresponding human PBMC bulk RNA-seq data available in Haemopedia. Second, higher resolution of distinct cell populations was obtained by single-cell RNA-sequencing (scRNA-seq) of PBMC. Seven PBMC samples were partitioned and sequenced that produced 28,810 single cell transcriptomes distributed across 36 clusters and classified into 13 general cell types including plasmacytoid dendritic cells (DC), conventional DCs, monocytes, B cell, conventional CD4 and CD8 αβ T cells, NK cells, and γδ T cells. Signature gene sets from the human Haemopedia data were assessed for relative enrichment in genes expressed in pig cells and integration of pig scRNA-seq with a public human scRNA-seq dataset provided further validation for similarity between human and pig data. The sorted porcine bulk RNAseq dataset informed classification of scRNA-seq PBMC populations; specifically, an integration of the datasets showed that the pig bulk RNAseq data helped define the CD4CD8 double-positive T cell populations in the scRNA-seq data. Overall, the data provides deep and well-validated transcriptomic data from sorted PBMC populations and the first single-cell transcriptomic data for porcine PBMCs. This resource will be invaluable for annotation of pig genes controlling immunogenetic traits as part of the porcine Functional Annotation of Animal Genomes (FAANG) project, as well as further study of, and development of new reagents for, porcine immunology.

## INTRODUCTION

A major goal of biological research is using genomic information to predict complex phenotypes of individuals or individual cells with specific genotypes. Predicting complex phenotypes is an important component of broad Genome-to-Phenome (G2P) understanding (Koltes et al., 2019), and investing in sequencing of multiple animal genomes, including pigs (*Sus scrofa*), for improved genome and cell functional annotation is key in solving the G2P question (Andersson et al., 2015; Giuffra et al., 2019). In addition to their major role in the world supply of dietary protein, pigs have anatomic, physiologic, and genetic similarities to humans and serve as biomedical models for human disease and regenerative medicine (reviewed in (Swindle et al., 2012; Kobayashi et al., 2018). Thus, deep annotation of porcine genome function would be a major milestone for addressing the G2P question. A highly contiguous porcine genome assembly with gene model-level annotation was recently published (Warr et al., 2020). However, this annotation is based primarily on RNA sequencing (RNA-seq) data from solid tissues, with few sample types representative of immune cells, with the exception of alveolar macrophages and dendritic cells (Auray et al 2016). Given the interaction of animal health and growth, any functional annotation of the porcine genome will be incomplete without deep analysis of expression patterns and regulatory elements controlling the immune system.

The transcriptomes of circulating immune cells serve as a window into porcine immune physiology and traits (Chaussabel et al., 2010; Mach et al., 2013; Schroyen and Tuggle, 2015; Auray et al., 2020). Blood RNA profiling has been used to understand variation in porcine immune responses (Huang et al., 2011; Arceo et al., 2013; Knetter et al., 2015; Munyaka et al., 2019) and genetic control of gene expression (Maroilley et al., 2017). One goal of such research is to develop gene signatures predictive of disease states (Berry et al., 2010) and predict responses to immunizations and/or infections (Chaussabel and Baldwin, 2014; Tsang et al., 2014), as has been demonstrated in humans. Whole blood is easily collected from live animals, but represents an extremely complex mixture of cell types. Estimates of gene expression in mixed samples are inherently inaccurate as cell composition differences are difficult to adjust for, complicating the interpretation of RNA differences across samples and treatments. Thus, starting from whole blood transcriptomic data, it is nearly impossible to link gene expression and regulation to a specific cell or cell type. To determine direct regulatory interactions, we must analyze specific cell populations and even individual cells. A cell type-specific understanding of peripheral immune cell gene expression patterns will thus enhance biological understanding of porcine immunity, reveal targets for phenotyping, and provide a comparison to other species.

Predominant immune cell populations in porcine peripheral blood mononuclear cell (PBMC) preparations are comprised mainly of monocytes, B-cells, and T-cells, with minor fractions of dendritic cells (DCs), natural killer (NK) cells, and NKT-cells also present. Porcine peripheral T-cell populations (reviewed in (Gerner et al., 2009a; Gerner et al., 2015) and DCs (Summerfield et al., 2015; Auray et al., 2016) are readily described based on phenotype, though deeper characterization of porcine immune cells could improve identification of valuable reagent targets and biological understanding of porcine immunity. T-cell populations are commonly grouped as αβ or γδ T-cells according to T-cell receptor (TCR) chain expression and further divided based on CD2, CD4, CD8α, and/or CD8β expression. Pigs have a unique CD2^-^ γδ T-cell lineage contributing to higher percentages of circulating γδ T-cells (Takamatsu et al., 2006) and unique αβ T-cells expressing both CD4 and CD8α (Zuckermann, 1999). Relatively little is known about different circulating B-cell populations in pigs, as reagents for phenotyping are limited.

Various technical approaches can be used to enrich or isolate specific cell populations, improving resolution of cell types for deeper interrogation of gene expression. Flow cytometry is used to characterize cells based on expression of cell type-specific protein markers, and live cells can be sorted by magnetic- and/or fluorescence-activated cell sorting (MACS/FACS) for use in subsequent assays. MACS/FACS enrichment followed by transcriptomic analysis can provide additional insight of gene expression in specific cell types, but cells expressing the same combination of markers are often still a heterogeneous mixture (Sutermaster and Darling, 2019). Some major subtypes of porcine immune cell populations can be labeled for cell sorting by existing antibody reagents (Gerner et al., 2009b), but some subtypes such as B-cells lack these resources.

An exciting alternative to sorting specific cell types for transcriptomic analysis is single-cell RNA-seq (scRNA-seq). Many scRNA-seq approaches do not require prior phenotypic/functional information or antibody reagents but instead rely on physical partitioning of cells to uniquely tagged transcripts from individual cells and sharpen resolution of subsequent transcriptomic analysis to single cells (Liu and Trapnell, 2016; Vieira Braga et al., 2016; Zheng et al., 2017). scRNA-seq methods have been applied to human PBMCs (Zheng et al., 2017) and provide more accurate and detailed analyses of transcriptional landscapes that can identify new cell types (Villani et al., 2017) when compared to other transcriptomic approaches. There are limitations to scRNA-seq, with tradeoffs in total genes detected per cell versus total cells captured for analysis, depending on the approach used (Wilson and Göttgens, 2018).

To deeply annotate the porcine genome for peripheral mononuclear immune cell gene expression and further inform phenotype and function of the heterogenous pool of immune cells in PBMC preparations, two approaches were used to isolate peripheral immune cells for RNA-seq. MACS followed by FACS was used to enrich for eight PBMC populations using population-specific cell surface markers, and RNA isolated from enriched populations was used for bulk RNA-seq (bulkRNA-seq) or a NanoString assay to evaluate gene expression. PBMCs were also subjected to droplet-based partitioning for scRNA-seq. Gene expression patterns of porcine immune cells using different approaches were compared to each other and to multiple human datasets. Complementary methods provided an improved annotation and deeper understanding of porcine PBMCs, as well as explicated datasets for further query by the research community.

## MATERIAL AND METHODS

### Animals and PBMC isolation

Four separate PBMC isolations were performed, with different animals used in each experiment. Cells were used for bulkRNA-seq, targeted RNA detection (NanoString), or scRNA-seq. PBMCs from experiments were used as follows: Experiment A (ExpA) for bulkRNA-seq of sorted populations from two ∼6-month-old pigs (A1, A2); Experiment B (Exp B) for NanoString and scRNA-seq from three ∼12-month-old pigs (B1, B2, B3); Experiment C (ExpC) for scRNA-seq from three ∼12-month-old pigs (C1, C2, C3); Experiment D (ExpD) for scRNA-seq from two ∼7-week-old pigs (D1, D2). All pigs were crossbred, predominantly Large White and Landrace heritage. All animal procedures were performed in compliance with and approval by NADC Animal Care and Use Committee. PBMCs were isolated, enumerated, and viability assessed as previously described (Byrne et al., 2020).

### Enrichment and sorting eight leukocyte populations by MACS/FACS

PBMCs were labeled with biotin labeled anti-porcine CD3ε (PPT3, Washington State University Monoclonal Antibody Center) for 15 min at 4 °C, mixing continuously. Cells were washed with Hank’s Balanced Salt Solution (HBSS), incubated with anti-biotin microbeads (Miltenyi Biotec), placed on LS columns, and separated into CD3ε^+^ and CD3ε^-^ fractions according to manufacturer’s directions (Miltenyi Biotec). CD3ε^+^ and CD3ε^-^ fractions were each fluorescently-sorted into four subpopulations based on surface marker expression shown in Figure 1 and Table 1. For NanoString assays, B-cells were sorted as CD3ε^-^CD172α^-^CD8α^-^; CD21 was not used for sorting. Each fraction for FACS was confirmed CD3ε^+^ or CD3ε^-^ by labeling with anti-mouse IgG1-PE-Cy7 to detect anti-CD3ε antibody used for MACS. Cells were sorted into supplemented HBSS using a BD FACSAria II with 70mm nozzle. After sorting, cells were pelleted and enumerated as described above. Sorted cell purity was >85% for each population. Cells were stained, sorted, and further processed within 10h of collection keeping cells on ice between processing steps.

**Figure 1.**
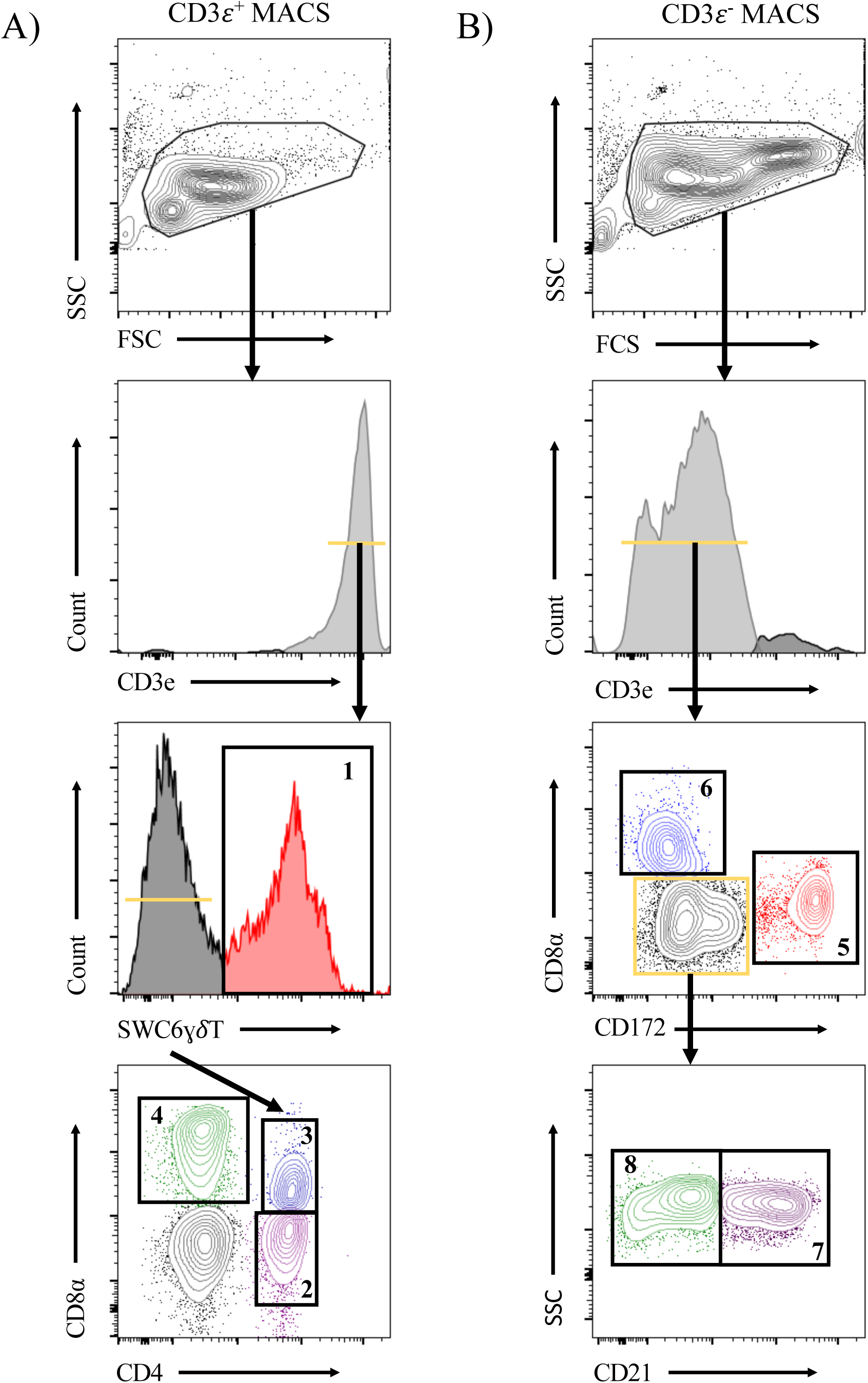
Representative plots for fluorescence-activated cell sorting (FACS) isolation of 8 leukocyte populations from pig peripheral blood mononuclear cells (PBMCs). Porcine PBMCs were first subjected to magnetic-activated cell sorting (MACS) to enrich for CD3ε+ and CD3ε- fractions. **A)** Cells in CD3ε^+^ MACS fraction were FACS gated on FSC vs SSC, doublets removed (not shown), and CD3ε^+^ cells were isolated into 4 population: SWC6^+^ γδ T-cells (gate 1), and the SWC6^-^ cells sorted as CD4^+^CD8α^-^ (gate 2), CD4^+^CD8α^+^ (gate 3), CD4^-^CD8α^+^ (gate 4) T-cells. **B)** Cells in CD3ε^-^ MACS fraction were FACS gated on FSC vs SSC, doublets removed (not shown), and CD3ε^-^ cells were isolated into 4 populations: CD172α^+^ myeloid lineage leukocytes (gate 5), CD8α^+^CD172^-^ NK cells (gate 6), and the remaining CD8α^-^ CD172α^-^, cells were isolated as CD21^+^ (gate 7) and CD21^-^ (gate 8) B-cells. Table 1 outlines abbreviations and sort criteria for each population.

**Table 1.**
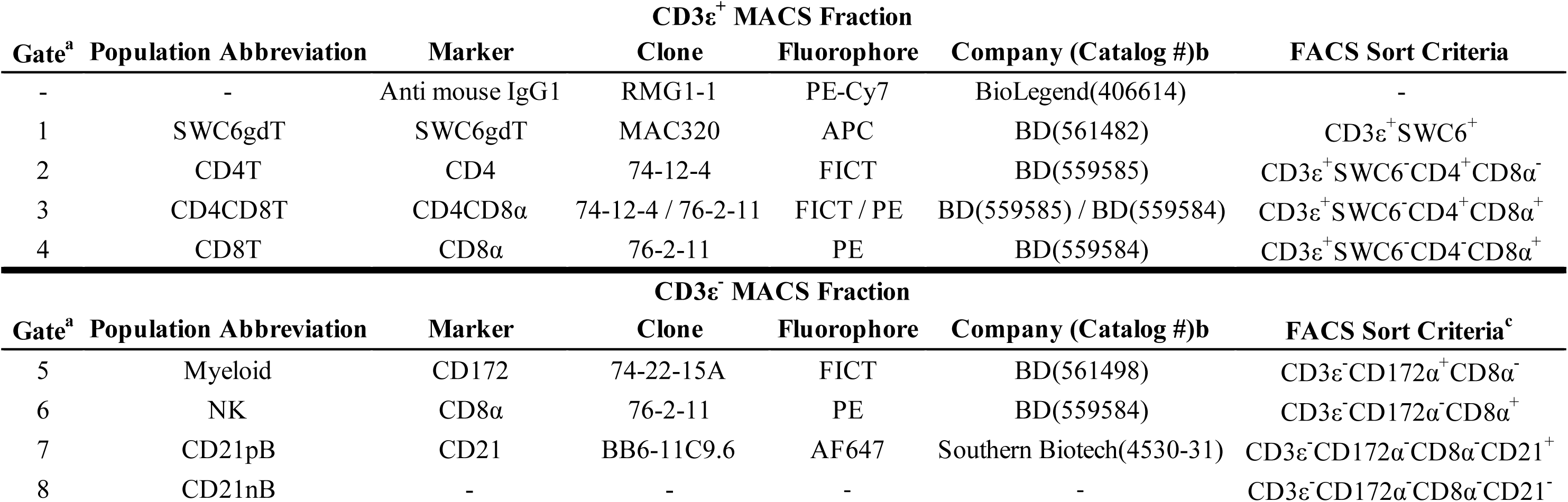
Abbreviations and phenotype information of pig sorted immune cells. ^a^ Refers to gate in Figure 1. ^b^ Reagents listed in materials and methods.

### RNA isolation for bulkRNA-seq/NanoString

BulkRNA-seq: after FACS, cells were pelleted, enumerated, and immediately lysed in RLT Plus buffer. RNA extractions were performed using the AllPrep DNA/RNA MiniKit (QIAGEN) following manufacturer’s instructions. Eluted RNA was treated with RNase-free DNase (QIAGEN). RNA quantity/integrity were assessed with an Agilent 2200 TapeStation system (Agilent Technologies). Samples used had RNA integrity numbers (RINs) ≥7.9. From ExpA, only one RNA sample for NK cells was used.

For NanoString assay: after FACS, cells were pelleted, enumerated, and immediately stored in Trizol. RNA extraction was performed using the Direct-zol RNA MicroPrep Kit (Zymo) with on-column DNase treatment following manufacturer’s instructions. RNA quantity and integrity were assessed as described above, with RINs ≥6.9. RNA was preserved at −80 °C until further use.

### BulkRNA-seq library preparation and data analysis

RNA was fragmented and 15 libraries prepared using the TruSeq Stranded Total RNA Sample Preparation Kit (Illumina). Libraries were diluted and pooled in approximately equimolar amounts. Pooled libraries were sequenced in paired-end mode (2×150-bp reads) using an Illumina NextSeq 500 (150 cycle kit).

#### Preprocessing, mapping, alignment, quality control

Data processing was performed as previously reported (Herrera-Uribe et al., 2020) using Sscrofa 11.1 genome and annotation v11.1.97 were used. Counts per gene of each sample in the two count tables were added together to get the final count table. Given that different types of immune cells have different transcriptome profiles (Hicks and Irizarry, 2015), YARN (Paulson et al., 2017), a tissue type-aware RNA-seq data normalization tool, was used to filter and normalize the count table. Genes with extremely low expression levels (<4 counts in at least one cell type) were filtered out using filterLowGenes(). The final count table contained 12,261 genes across 15 samples, which was then normalized using normalizeTissueAware(), which leverages the smooth quantile normalization method (Hicks et al., 2018).

Data quality control was performed using DESeq2 (v1.24.0) (Love et al., 2014) within RStudio s (v1.2.1335). Regularized log-transformation was applied to the normalized count table with the rld function. Then principal component analysis (PCA) and sample similarity analyses were carried out and visualized using plotPCA() and distancePlot(), respectively. Heatmaps to display enriched genes were created using pheatmap (v1.0.12) within RStudio.

#### Cell type-enriched and cell type-specific gene identification

The normalized count table was used for differential gene expression (DGE) analysis with DESeq2 by setting the size factor for each sample to 1. A generalized linear model was fitted for each gene in the count table, with negative binomial response and log link function of the effect of cell types and pig subjects. nbinomWaldTest() was used to estimate and test the significance of regression coefficients with the following explicit parameter settings: betaPrior=FALSE,maxit=50000,useOptim=TRUE,useT=FALSE,useQR=TRUE. Cell type-enriched genes and cell type-specific genes were identified using the results function separately. A gene was labeled as <u>cell-type enriched</u> if the expression level (averaged across replicates) in one cell type was at least 2x higher than the average across all remaining cell types and adjusted p-value <0.05. A gene was labeled as <u>cell type-specific</u> if the averaged expression level in one cell type was at least 2x higher in pairwise comparison to the average in each other cell type and adjusted p-value <0.05 (Benjamini and Hochberg, 1995). Heatmaps to display specific genes were created as mentioned above.

For cross-species comparison, human hematopoietic cell (Haemopedia) RNA-seq expression data (Hilton Laboratory at the Walter and Eliza Hall Institute^1^) was used. Only orthologous genes with one-to-one matches between human and pig (orthologous gene list obtained from BioMart (Durinck et al., 2009) were compared. Orthologous gene transcript per million (TPM) values from naive and memory B-cells, myeloid dendritic cells (myDC), myeloid dendritic cells CD123+ (CD123PmDC), plasmacytoid DC (pDC), monocytes, NK cells, CD4T and CD8T cells from healthy donors were used (Choi et al., 2019). Spearman rank correlation analyses was performed to identify correlation between orthologous gene expression levels (absolute TPM) in pig and human sorted populations. Significance level was set at P<0.05 and level of Spearman’s rank correlation coefficient (rho) was defined as low (<0.29), moderate (0.3-0.49), and strong (0.5-1) correlation.

#### Gene Ontology (GO) enrichment analysis

Metascape analysis (Zhou et al., 2019) was performed for GO analysis of the top 25% enriched genes and specific genes identified as described above, with threshold p-value <0.01. Several terms were clustered into the most enriched GO term. Term pairs with Kappa similarity score >0.3 were displayed as a network to show relationship among enriched terms. Terms associated with more genes tended to have lower P-values. All networks displayed were visualized using Cytoscape. All Ensembl Gene IDs with detectable expression level in each cell type were used as the background reference.

### NanoString assay and data analysis

A total of 230 test genes with nine housekeeping genes, eight positive and nine negative control genes were chosen for gene expression quantification on the NanoString nCounter analysis system (NanoString Technologies) using custom-made probes. The custom designed CodeSet was selected from genes and pathways associated with porcine blood, lung, lymph node, endometrium, placenta or macrophage response to infection with a porcine virus (Van Goor et al., 2020). RNA samples were diluted to 25-100 ng/ul in RNase-free water, and 5 ul of each sample was used in the assay using manufacturer’s instructions with the nCounter Master kit.

The nCounter analysis system produces discrete count data for each gene assayed within each sample. We used the NanoString software nSolver Analysis Software (v3.0, NanoString Technologies), following manufacturer’s instructions. The nSolver corrected for background based on negative control samples, performed within-sample normalization based on positive control probes, and performed normalization across samples using the median expression values of housekeeping genes (*GAPDH, HMBS, HPRT1, RPL32, RPL4, SDHA, TBP, TOP2B, YWHAZ*), providing confidence in our normalization method.

All statistical analyses were performed using the statistical programming language R v3.5. Raw count data were normalized using normalizationFactors() and NanoStringDataNormalization() from NanoStringDiff (v1.1.2.0) (Wang et al., 2016). One gene (*ISG20*) without detected expression in any samples was removed. Hierarchical clustering and PCA suggested there were substantial hidden variations among the expression data. Surrogate variable analysis has been shown to be a powerful method to detect and adjust for hidden variations in high throughput gene expression data (Li et al., 2014; Qian Liu, 2016), so surrogate variable analysis was applied to remove further hidden variations in the gene expression data using svaseq() from sva (v3.30.1) (Leek et al., 2012). A full model with cell subpopulations and RINs as independent variables, and a reduced model with RINs as the only independent variable were used. Three surrogate variables were estimated and used to adjust for the hidden variations.

Gene expression values were transformed to log_2_(TPM) using voom() from limma (Law et al., 2014). Linear mixed effect models were used to fit the transformed gene expression data by using lmer()in lme4 (Bates et al., 2015). The model included fixed effect for cell subpopulation, RIN, the three surrogate variables, and random effect for each animal. One minus Spearman correlation coefficient was used as distance measure for gene clustering, and Euclidian distances was used for sample clustering.

Additionally, Spearman correlation analysis was performed to assess the correlation between bulkRNA-seq and NanoString results. The significant level was set at P <0.05, and the level of Spearman’s rank correlation coefficient (rho) was defined as described above.

### scRNA-seq library preparation

PBMC isolation experiments were performed at different times and samples sequenced in different runs. For ExpB, 1×10^7^ viable PBMCs per animal were cryopreserved according to 10X Genomics Sample Preparation Demonstrated Protocol, shipped on dry ice to University of Minnesota’s Core Sequencing Facility, and thawed, partitioned, and scRNA-seq libraries prepared. For ExpC/ExpD, freshly isolated PBMCs were transported on ice to Iowa State University Core Sequencing Facility for partitioning and library preparation. Partitioning and library preparation were performed according to Chromium Single Cell 3’ Reagent Kits v2 User Guide (10X Genomics). For all experiments, 100 base paired-end reads were sequenced on an Illumina HiSeq3000 at ISU Core Sequencing Facility. One sample from ExpB was omitted from further analyses due to poor sequence performance.

### scRNA-seq data analysis

#### Read alignment/gene quantification

Raw read quality was checked with FASTQC^1^. Reads 2 (R2) were corrected for errors using Rcorrector (Song and Florea, 2015), and 3’ polyA tails >10 bases were trimmed. After trimming, R2 >25 bases were re-paired using BBMap^2^. *Sus scrofa* genome Sscrofa 11.1 and annotation GTF (v11.1.97) from Ensembl were used to build the reference genome index (Yates et al., 2020). The annotation file was modified to include both gene symbol (if available) and Ensembl ID as gene reference (e.g. GZMA_ENSSSCG00000016903) using custom Perl scripts. Processed paired-end reads were aligned and gene expression count matrices generated using CellRanger (v4.0; 10X Genomics) with default parameters. Only reads that were confidently mapped (MAPQ=255), non-PCR duplicates with valid barcodes, and unique molecular identifiers (UMIs) were used to generate gene expression count matrices. Reads with same cell barcodes, same UMIs, and/or mapped to the same gene feature were collapsed into a single read.

#### Quality control/filtering

CellRanger output files were used to remove ambient RNA from each sample with SoupX (Young and Behjati, 2020) function autoEstCont(). Corrected non-integer gene count matrices were outputted in CellRanger file format using DropletUtils (Lun et al., 2019) function write10xCounts() and used for further analyses. Non-expressed genes (sum zero across all samples) and poor quality cells (>10% mitochondrial genes, <500 genes, or <1,000 UMIs per cell) were removed using custom R scripts and Seurat (Stuart et al., 2019). Filtered count matrices were generated using write10xCounts() and used for further analyses. High probability doublets were removed using Scrublet (Wolock et al., 2019), specifying 0.07 expected doublet rate and doublet score threshold of 0.25.

#### Integration, visualization, and clustering

Post-quality control/filtering gene counts/cells from each sample were loaded into a Seurat object and transformed individually using SCTransform(). Data were integrated with SelectIntegrationFeatures(), PrepSCTIntegration(), FindIntegrationAnchors(), and IntegrateData() with default parameters. PCA was conducted with RunPCA(), and the first 14 principal componenets (PCs) were selected as significant based on <0.1% variation of successive PCs. Significant PCs were used to generate two-dimensional t-distributed stochastic neighbor embedding (t-SNE) and uniform manifold approximation and projection (UMAP) coordinates for visualization with RunTSNE() and RunUMAP(), respectively, identify nearest neighbors and clusters with FindNeighbors() and FindClusters() (clustering resolution = 1.85), respectively, and perform hierarchical clustering with BuildClusterTree(). Counts in the RNA assay were further normalized and scaled using NormalizeData() and ScaleData().

#### Differential Gene Expression (DGE) analyses

Normalized counts from the RNA assay were used for DGE analyses. Differentially-expressed genes (DEGs) between pairwise cluster combinations were calculated using FindMarkers(). DEGs in one cluster relative to the average of all other cells in the dataset were calculated using FindAllMarkers(). The default Wilcoxon Rank Sum test was used for DGE analyses. Genes expressed in >20% of cells within one of the cell populations being compared, with |logFC|>0.25, and adjusted p-value <0.05 were considered DEGs.

#### Gene set enrichment analyses (GSEA)

Enrichment of gene sets within our porcine scRNA-seq dataset were performed using AUCell (v1.10.0) (Aibar et al., 2017). Enriched genes in sorted porcine bulkRNA-seq populations were identified as described in preceding methods. Log2FC values were used to curate gene sets of genes enriched in the top 25%, 20%, 15%, 10%, 5%, or 1% of bulkRNA-seq populations. Gene sets from human bulkRNA-seq cell populations (Choi et al., 2019) were recovered by performing a High Expression Search on the Haemosphere website^1^, setting Dataset=Haemopedia-Human-RNASeq and Sample group=celltype. Gene sets for CD4:+ T-cell; CD8:+ T-cell; Memory B-cell; Monocyte; Myeloid Dendritic Cell; Myeloid Dendritic Cell CD123+; Naïve B-cell; Natural Killer Cell; and Plasmacytoid Dendritic Cell options corresponded to CD4T, CD8T, MemoryB, Monocyte, mDC, CD123PmDC, NaïveB, NK, and pDC designations, respectively. Genes with high expression scores >0.5 (lower enrichment level) or >1.0 (higher enrichment level) were selected and filtered to include only one-to-one gene orthologs as described in preceding methods. Human gene identifiers were converted to corresponding porcine gene identifiers or gene names used for scRNA-seq analyses.

Within each cell of the finalized scRNA-seq dataset, gene expression was ranked from raw gene counts. Area under the curve (AUC) scores were calculated from the top 5% of expressed genes in a cell and the generated gene sets. Higher AUC scores indicated a higher percentage of genes from a gene set were found amongst the top expressed genes for a cell. For overlay of AUC scores onto UMAP coordinates of the scRNA-seq dataset, a threshold value was manually set for each gene set based on AUC score distributions. For visualization by heatmap, AUC scores were calculated for each cell, scaled relative to all other cells in the dataset, and average scaled AUC scores were calculated for each cluster.

#### Deconvolution analysis (CIBERSORTx)

To deconvolve cluster-specific cell subsets from bulkRNA-seq of sorted populations, CIBERSORTx (Newman et al., 2019) was used to derive a signature matrix from scRNA-seq data. 114 cells were taken from each cluster using the Seurat subset() function and labelled with corresponding cluster identities. Cluster-labeled cells were used to obtain a single-cell reference matrix (scREF-matrix) that was used as input and run on CIBERSORTx online server using “Custom” option. Default values for replicates (5), sampling (0.5), and fraction (0.0) were used. Additional options for kappa (999), q-value (0.01), and No. Barcode Genes (300-500) were kept at default values. CIBERSORTx scREF-matrix was used to impute cell fractions from the bulkRNA-seq of sorted cell population “mixtures”. The mixture file (TPM values) was used as an input and run on CIBERSORTx online server using the “Impute Cell Fractions” analysis with the “Custom” option selected, and S-mode batch-correction was applied. Cell fractions were run in relative mode to normalize results to 100%. The number of permutations to test for significance were kept at default (100). Resulting output provided estimated percentages of what scRNA-seq clusters defined each bulkRNA-seq sorted cell population.

#### Reference-based label transfer/mapping and de novo integration/visualization

A CITE-seq dataset of human PBMCs (Hao et al., 2020) was used to transfer cell type annotations onto our porcine scRNA-seq dataset. Due to the cross-species comparison, we distilled human reference and pig query datasets to only include 1:1 orthologous gene, and human reference dataset was re-normalized and integrated mirroring previous methods (Hao et al., 2020). Each sample of the porcine query dataset was separately normalized using SCTransform. Anchors were found between the human reference and each pig query sample using FindTransferAnchors. Identified anchors were used to calculate mapping scores for each cell using MappingScore. The mapping scores provided a 0-1 confidence value of how well a porcine cell was represented by the human reference dataset. Prediction scores were calculated using available level 2 cell types from the human reference dataset. Prediction scores provided a 0-1 percentage value for an individual cell type prediction, based on how many nearby human cells shared the same cell type annotation that was predicted. Predicted cell annotations were projected back onto original UMAP of the porcine dataset. Cluster-averaged prediction and mapping scores were also calculated.

In order to identify cells from the porcine dataset that were not well represented by the human reference dataset the two datasets were integrated to perform *de novo* visualization by merging the two datasets and their respective sPCAs to create a new UMAP. From two-dimensional *de novo* UMAP, porcine cells that did not overlap with human cells were identified.

#### Cluster subsetting

For deeper analyses of only subsets of clusters in the scRNA-seq dataset, cells belonging to only selected clusters were place in a new Seurat object using subset(). Genes with zero overall expression in the new data subset were removed using DietSeurat(), and counts were re-scaled with ScaleData(). Original cluster designations and PCs were left intact. UMAP/t-SNE visualization, hierarchical clustering, and DGE analyses were re-performed as described in the original analyses. Pairwise DGE analyses were not re-performed.

#### Random Forest (RF) Modeling

The RF models provided an estimate of cluster similarity based on error rates. The R packages caret^1^ and ranger^2^ were used to create RF models trained on cluster identities of cells. A normalized count matrix was used as input data for RF models. Each cell was labeled by its previously defined cluster. Two different types of models were created: (1) pairwise models where training data included only cells from two different clusters (ex. Clusters 0 & 3); (2) models where training data included cells from all clusters of a specified dataset (ex. all γδ T cell clusters). Each model was trained on the cluster identity of each cell, with trees created=500, target node size = 1, variables=14,386, variables to sample at each split (Mtry)=119. Each tree in the model is grown from a bootstrap resampling process that calculates an out-of-bag (OOB) error that provides an efficient and reasonable approximation of the test error. Variable importance was used to find genes or sets of genes that can be used to identify certain types of cells or discriminate groups of cells from one another. RF models are advantageous because they can provide ranked lists of genes most important for discriminating cells between different clusters. This method was used to identify groups of important genes to supplement single DGE analyses. Variable importance was assigned by measuring node impurity (Impurity) and using permutations (Permutation). Features that reduced error in predictive accuracy are ranked as more important. High error rate in the model suggests cells from the groups being compared are more similar to each other, whereas low error rate suggests cells from each cluster are unique.

#### Gene name replacement

Several gene names/Ensembl IDs used for data analysis were replaced in main text/figures for the following reasons: gene symbol was not available in the annotation file but was available under the gene description on Ensembl, gene symbol was updated in future Ensembl releases, or multiple Ensembl IDs corresponded to a single gene symbol. Affected genes included: ABI3=ENSSSCG0000003522, ABRACL=ENSSSCG00000004145, AP3S1=ENSSSCG00000037595, CCDC12=ENSSSCG00000011329, CCL23=ENSSSCG00000033457, CD163L1=ENSSSCG00000034914, CDNF=ENSSSCG00000039658, CR2=ENSSSCG00000028674, CRIP1=ENSSSCG00000037142, CRK=ENSSSCG00000038989, EEF1A1=ENSSSCG00000004489, FCGR3A=ENSSSCG00000036618, GBP1=ENSSSCG00000024973, GBP7=ENSSSCG00000006919, GIMAP4=ENSSSCG00000027826, GZMA=ENSSSCG00000016903, HMGB1=ENSSSCG00000009327, HOPX=ENSSSCG00000008898, IFITM1=ENSSSCG00000014565, IGLL5=ENSSSCG00000010077, KLRB1B=ENSSSCG00000034555, KLRC1=ENSSSCG00000000640, KLRD1=ENSSSCG00000026217, MAGOHB=ENSSSCG00000000635, MAL=ENSSSCG00000040098, MAN2B1=ENSSSCG00000013720, MDK=ENSSSCG00000013260, MYL12A=ENSSSCG00000003691, NT5C3A=ENSSSCG00000022912, PRKCH=ENSSSCG00000005095, PTTG1=ENSSSCG00000017032, RPL14=ENSSSCG00000011272, RPL22L1=ENSSSCG00000036114, RPL23A=ENSSSCG00000035080, RPL35A=ENSSSCG00000040273, RPS15A=ENSSSCG00000035768, RPS19=ENSSSCG00000003042, RPS27A=ENSSSCG00000034617, RPS3=ENSSSCG00000014855, RPS8=ENSSSCG00000003930, S100B=ENSSSCG00000026140, SIRPA=ENSSSCG00000028461, SLA-DQA1=ENSSSCG00000001456, SLA-DRA=ENSSSCG00000001453 (listed as HLA-DRA in the gene annotation used), SLA-DRB1=ENSSSCG00000001455, SLPI=ENSSSCG00000022258, SPIB=ENSSSCG00000034211, TMSB4X=ENSSSCG00000012119, TXN=ENSSSCG00000005453, WIPF1=ENSSSCG00000027348.

## RESULTS

### BulkRNA-seq revealed common and distinct transcriptomes in circulating immune cells

Eight immune cell populations (Table 1) were sorted by cell-surface marker phenotypes for transcriptomic profiling by bulkRNA-seq (Figure 1) using primarily criteria previously outlined (Gerner et al., 2009b), with some modifications. Our protocol utilized an antibody reactive to swine workshop cluster 6 (SWC6) protein to identify γδ T-cells, but the antibody only labels CD2^-^ γδ T-cells (Yang and Parkhouse, 1996; Davis et al., 1998; Stepanova and Sinkora, 2013; Sedlak et al., 2014). CD2^+^ γδ T-cells were likely sorted into the CD3ε^+^CD4^-^CD8α^-^ fraction that was not retained or the CD8T (CD3ε^+^CD4^-^CD8α^+^) population (Davis et al., 1998; Stepanova and Sinkora, 2013; Sedlak et al., 2014). A pan-B-cell marker for pigs is not currently available, so B-cells are often characterized through a series of negative gates. Cells in the CD3ε^-^ fraction were considered B-cells if they also lacked expression of CD172α and CD8α. B-cells characterized in this manner were further terminally sorted into B-cell populations with or without CD21 (complement receptor 2) expression (CD21pB and CD21nB, respectively; Figure 1, gates 7 and 8 respectively). We acknowledge that the CD21nB gate likely contained other circulating cell types that were not sorted through positive gating approaches.

Transcriptomic profiles of sorted cell populations were constructed by bulk RNA-seq, and relationships among porcine immune cell transcriptomes were assessed and visualized through dimensionality reduction and hierarchical clustering (Figure 2A and 2B and Supplementary File 1). Specifically, T-cell populations (SWC6gdT, CD4T, CD4CD8T, CD8T), B-cell populations (CD21pB, CD21nB), myeloid leukocyte populations (Myeloid), and a single NK cell population (NK) were well separated from each other (Figure 2A) by PCA. Replicates of specific sorted cell populations clustered most closely together, while within T-cell populations or B-cell populations, considerable transcriptional similarity was observed (Figure 2B).

**Figure 2.**
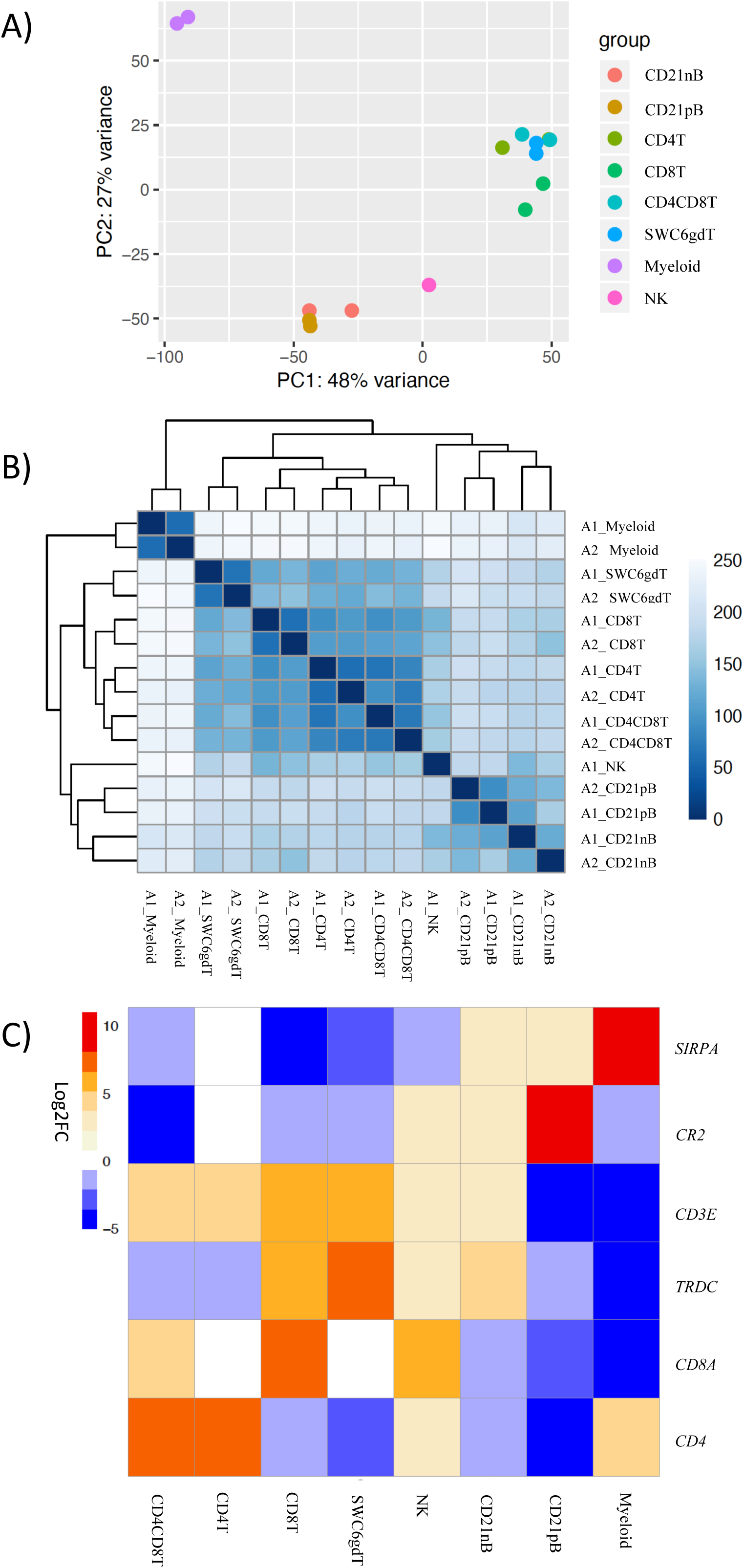
Transcriptional expression patterns of immune cells are distinct and cluster more by progenitors. **A)** Principal component analysis of transformed RNA-seq reads counts for whole transcriptomes. Axis indicate component scores. **B)** Heat map depicting hierarchical clustering of sample-to-sample distance. Gene expression for whole transcriptomes were used to calculate sample to sample Euclidean distance (color scale) for hierarchical clustering. **C)** Heatmap showing cell-type enriched gene values (Log2FC) between sorted immune cells. Gene coding proteins that were used for cell sorting were display.

The total number of expressed genes in each sorted population was similar (Supplementary File 1). Significantly enriched genes (SEGs) with expression significantly different and at least 2x greater than the average of all other cell populations (see Methods) were identified for each sorted population (Supplementary File 2). Notably, around 12-18% of SEGs are not fully annotated (no symbol/gene name) in the Sscrofa 11.1 genome and annotation v11.1.97. The SWC6gdT population had the highest number of SEGs (3,591), while the NK population had the fewest (1,885) (Table 2). SEG lists were queried for corresponding protein targets used to sort cells, if known, to confirm enrichment of expression of genes corresponding to protein phenotypes (Figure 2C). Expression of *SIRPA*^*^ (encoding CD172α) had the highest fold-change in the Myeloid population, and *CR2 (*encoding CD21*, ENSSSCG00000028674*), was highest in the CD21pB population, as would be predicted based on protein phenotypes. The two CD4^+^ T-cell populations (CD4T and CD4CD8T) had the highest fold-change for *CD4.* The CD8T population had the highest fold-change for *CD8A*, with CD4CD8T and NK populations also having near a log_2_FC enrichment value of 5, in line with these populations also expressing CD8α. The SWC6gdT population had the highest fold-change for *TRDC*, though CD8T and CD21nB populations also had enrichment for *TRDC*. As noted previously, it’s unlikely our sorting for γδ T-cells based on SWC6 captured all γδ T-cells, thus some γδ T-cells may be represented in other sorted populations. Thus, the CD8T population is likely comprised not only CD8α^+^ αβ T-cells, but also potentially SWC6^-^ γδ T-cells expressing CD8α.

**Table 2.**
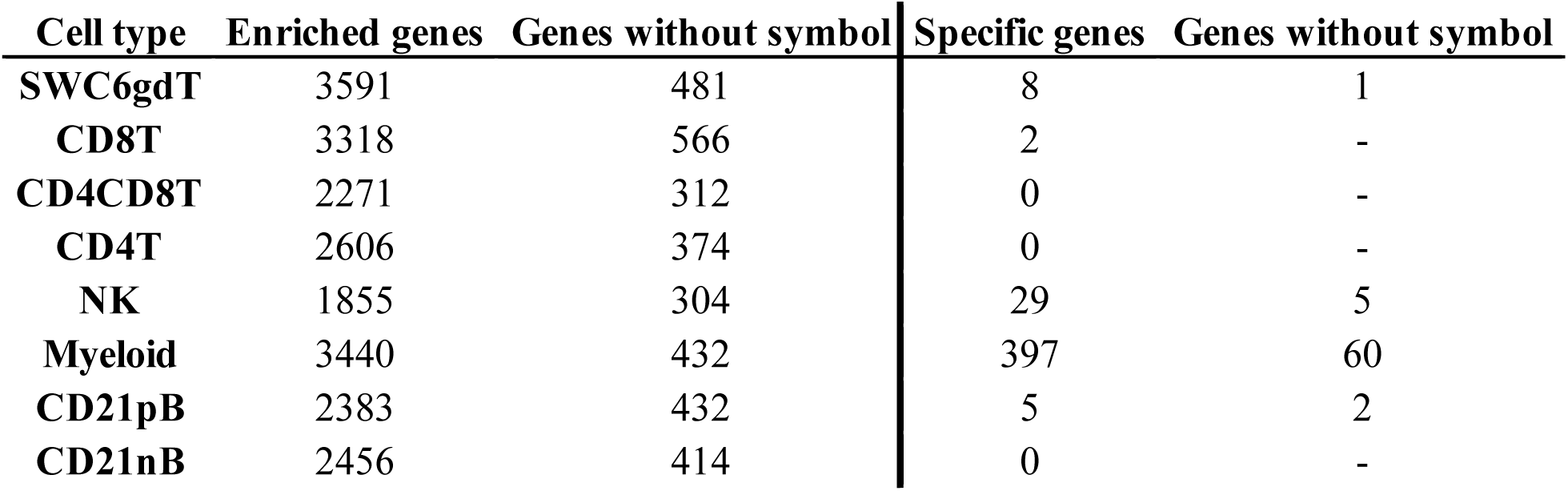
Cell type-enriched and cell type specific genes identified in pig sorted immune cells.

A subset of SEGs (25% highest log2FC values) for each sorted population, referred to as highly enriched genes (HEGs) that distinguish different circulating pig immune cell populations, were used for data visualization and GO analysis. The log_2_FC values for HEGs were clustered and visualized in Figure 3 (four CD3ε^-^ populations) and Supplementary Figure 2 (four CD3ε^+^ populations). GO analyses using HEG lists for each cell population indicated enrichment for biological processes characteristic of each respective cell population, depicted as networks of similar terms (Figure 3E-3H, Supplementary File 3, Supplementary Figure. 2E-2H). Terms for Myeloid HEGs included Myeloid leukocyte activation and response to bacterium (Figure 3E), and terms for NK HEGs included positive regulation of cell killing and natural killer cell mediated cytotoxicity (Figure 3F). Many terms enriched for CD21pB HEGs overlapped with those for CD21nB HEGs, as 38% of HEGs were shared between these populations (Figure 3C, 3D). Thus, top GO terms for B-cells, including adaptive immune response and B-cell proliferation were present in both populations (Figure 3G and 3D). However, some GO terms were unique to either B-cell population. GO related to B-cell activation, such as positive regulation of B-cell activation/proliferation processes associated with B-cell receptor signaling, were identified exclusively for CD21pB HEGs. For CD21nB HEGs, processes associated with humoral immunity and red blood cell processes such as coagulation or platelet activation were noted, which could indicate contamination of different cell types given the non-specific cell sorting approach used for CD21nB cells (Figure 1). For all sorted T-cell populations (CD8T, CD4T, CD4CD8T and SWC6gdT), HEG lists showed overlap (Supplementary Figure 2A-D). GO terms included T-cell activation, T-cell receptor signaling pathway, cytokine-cytokine receptor interaction and biological processes related to cytotoxicity activity (Supplementary Figure 2E-2F). Overall, GO exploration of HEGs for sorted populations provided evidence that sorted immune cells represented expected immune cell functions.

**Figure 3.**
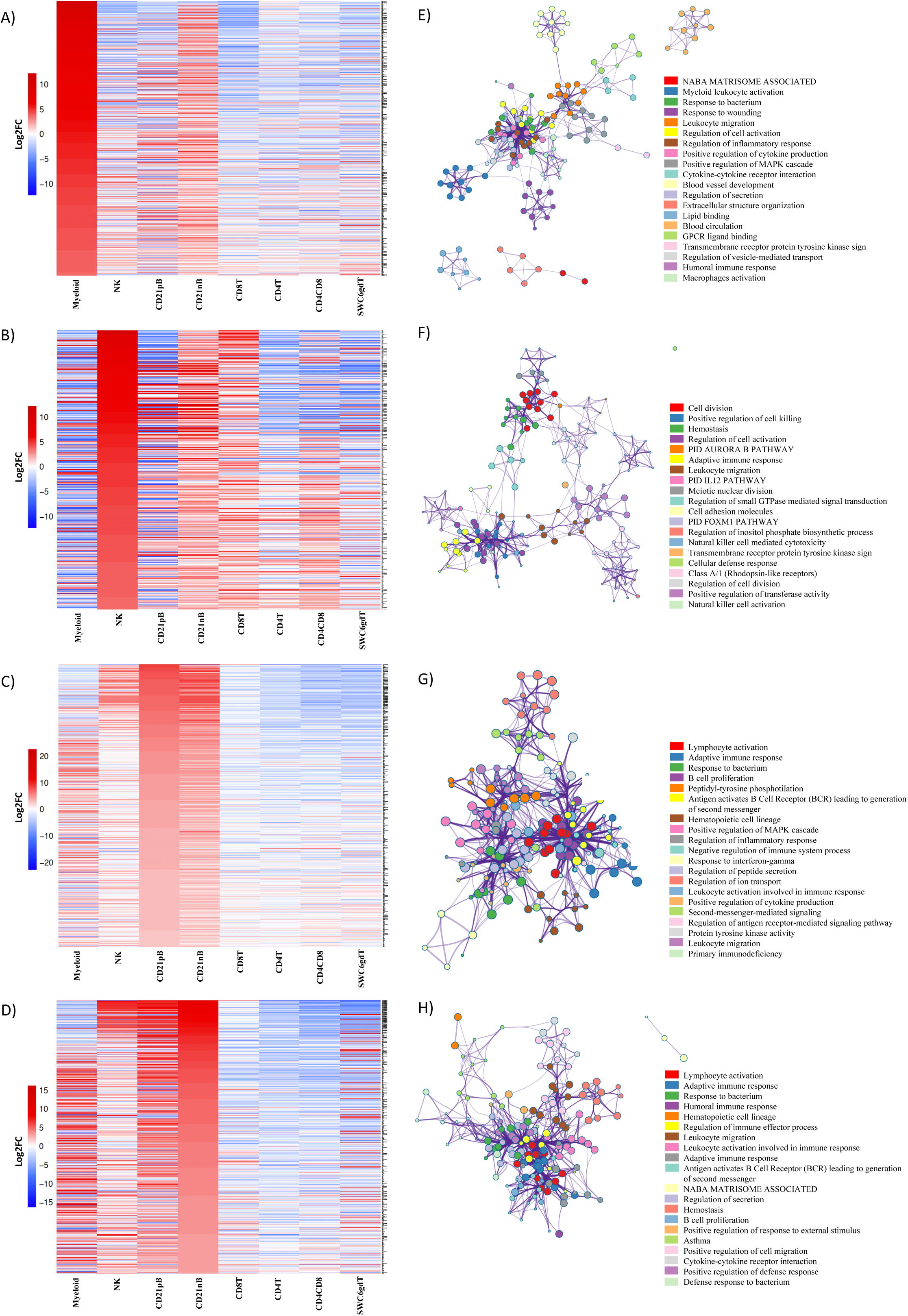
Top 25% highly enriched genes in CD3-sorted cells. Heatmap showing in decreasing order the top 25% of highly enriched genes in **A**) myeloid, **B**) NK, **C**) CD21pB and **D)** CD21nB-cells. Ontology enrichment clusters of the top 25% highly enriched genes of **E**) myeloid, **F**) NK, **G**) CD21pB and **H)** CD21nB-cells. The most statistically significant term within similar term cluster was chosen to represent the cluster. Term color is given by cluster ID and the size of the terms is given by –log10 P-value. The stronger the similarity among terms, the thicker the edges between them.

The TPM values of expressed genes in sorted porcine cells were compared with orthologous human genes expressed in sorted human naïve hematopoietic cells from the Haemopedia (Choi et al., 2019) in order to identify cell-specific transcriptome similarities across species. Gene expression correlations assessed by Spearman’s rank correlation indicated highly significant and moderately strong correlations (rho=0.30-0.43, P<2.2e-16) between porcine and anticipated corresponding human immune cell populations (Supplementary Figure 3, Supplementary File 4). A closer evaluation of genes reported as canonical cell markers for different mouse and human peripheral immune cell populations^1^ and expression of those genes in each of the sorted porcine populations revealed several commonalities. Specifically, genes such as *EBF1*, *CD19*, *MS4A1, CD79B*, *PAX5*, *HLA-DOB* (in CD21nB, CD21pB); *CD28* (in CD8T, CD4T, CD4CD8T); *CD5* (in CD8T, CD4T, CD4CD8T, SWC6gdT); *GZMA*, *GNLY*, *CCL5*, *KLRK1*, *KLRB1*, *CD244* (in NK, CD8T); and *VLDLR*, *NLRP3*, *CD14*, *STEAP4*, *CD163*, *DEFB1* (in Myeloid) for human cells showed specific enrichment in respective porcine populations (Supplementary Figure 4). Thus, additional query confirmed sorted porcine immune cell populations were equivalent to human counterparts in many ways.

### High homogeneity amongst sorted T-cell and B-cell populations and transcriptomic distinctions in Myeloid and NK populations

Pairwise DGE analyses between the cell populations identified genes with transcript abundance at least 2x higher in one population than in all other populations (adjusted p-value <0.05, see Methods) which we define as cell type-specific. Consistent with PCA (Figure 2A), more cell type-specific genes were identified in the Myeloid population than in NK, T or B-cells. In total, we identified 2, 5, 8, 29, and 397 cell type-specific genes for CD8T, CD21pB, SWC6gdT, NK, and Myeloid populations, respectively (Table 2, Supplementary Figure 5). GO analyses using cell type-specific genes for the Myeloid population resulted in enrichment of terms such as Myeloid leukocyte, cytokine-cytokine receptor interaction, and pattern recognition receptor activity (Supplementary Figure 5, Supplementary file 3). Next, we determined if the cell type-specific genes identified were present in the list of HEGs for each population. In total, 2, 2, 5, 14, and 271 cell type-specific genes were identified in respective HEG lists for CD8T, CD21pB, SWC6gdT, NK, and Myeloid populations, respectively (Table 3), indicating the most highly-enriched cell type-specific genes were present in NK and Myeloid populations. Cell type-specific genes could not be identified for the remaining three sorted populations (CD4T, CD4CD8T, and CD21nB) using the criteria described above, indicating between-population transcriptional heterogeneity even for these enriched populations.

**Table 3.**
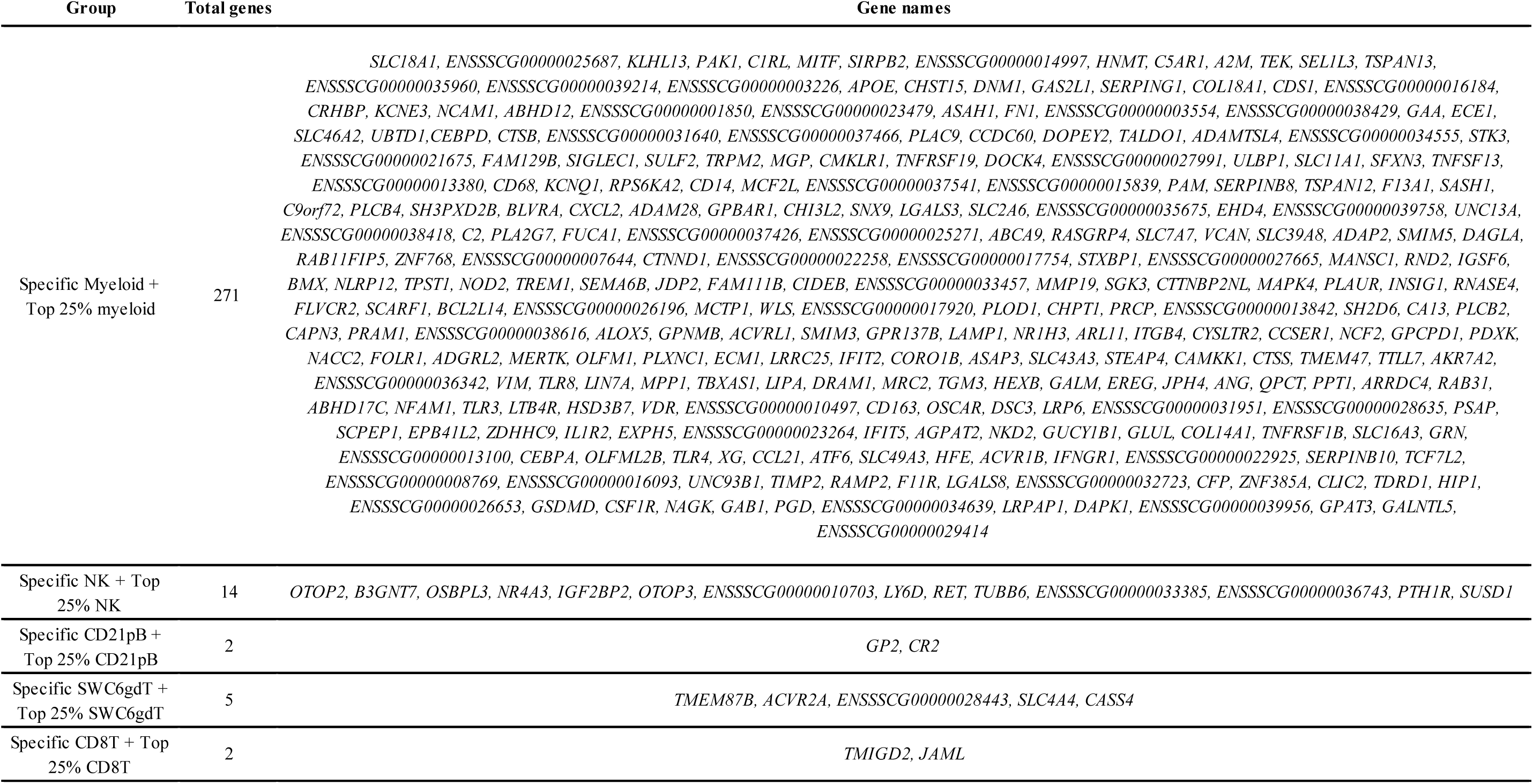
Specific highly enriched genes in myeloid, NK, CD21pB, SWC6gdT and CD4CD8T-cells.

We then explored immune cell transcriptomic patterns to identify genes that could expand our knowledge of pathways active in specific cell populations, as well as predict new genes suitable to use for molecular analyses in immunology studies. Of interest, we found a remarkably high number of HEGs in our Myeloid population (Table 3), including immune-related genes involved in TLR signaling (*CD14*, *CD36*, *TLR2/3/4/8/9*, *NOD2*) and cytokine activity (*CSF1R*, *CSF2RA*, *CSF3R*, *IFNGR1*, *IL1B*, *IL1RAP*, *CXCR2*, *CCL21*, *CCL23*, *TNFRSF1B*, *IL1R2*, *TNFSF13*, *TNFSF13B*, *TNFRSF21*, *CXCL16*, *CCR2*). In NK cells fewer specific genes were detected than the Myeloid population (Table 3), with genes such as *OTOP2*, *OTOP3*, *OSPBL3*, *LY6D*, *RET* related to cytotoxic activity, a typical characteristic of NK cells (Rusmini et al., 2013; Rusmini et al., 2014; Belizário et al., 2018; Costanzo et al., 2018; Tu et al., 2018; Upadhyay, 2019), although their function in porcine NK cells is unexplored. In CD21pB cells, the gene for CD21 (*CR2*) used for sorting the B-cell populations was predicted to be a HEG. The SWC6gdT population showed specific expression of *AVCR2A*, which is a Th17 cell specific gene in mice (Ihn et al., 2011) and regulates the proliferation of γδ T-cells in murine skin (Antsiferova et al., 2011). The CD8T population specifically expressed *TMIGD2* (a CD28 family member) and *JAML*, which encode T-cell transmembrane proteins (Zhu et al., 2013; Alvarez et al., 2015; Krueger et al., 2017).

Finally, we compared pair-wise transcriptome differences between our porcine sorted CD4T and CD8T populations (Supplementary File 2) with the comparable populations from a previous study (Foissac et al., 2019). Even though the sorting approaches were different, 85% of the genes more highly expressed in in CD4T compared to CD8T, respectively, were detected by Foissac and colleagues in their respective CD4^+^ to CD8^+^ comparison. Similar overlap was found (87%) for the genes more abundant in the “CD8+ high” list, while little overlap was found in the inverse comparisons (2.5% and 1%), strongly indicating these cell type gene expression patterns were similar between studies. However, given the lack of identification of cell-type specific genes for CD4T and CD8T populations, shared gene expression patterns may not be surprising.

### NanoString assay validated bulkRNA-seq

RNA abundance of each gene target (Supplementary File 5) in each sample was used to perform a hierarchical clustering analysis (Supplementary Figure 6). Similar to relationships observed in the bulkRNA-seq dataset, biological replicates clustered most closely together. T-cell populations (SWC6gdT, CD4T, CD4CD8T, CD8T) were more similar to each other than to other populations, with the exception of NK cells. RNA abundance for the genes encoding the marker proteins used for sorting cell populations confirmed cell identity in NanoString assays (Supplementary Figure 7). RNA abundance for each tested gene and cell population is included in Supplementary File 5. To validate gene expression levels calculated by bulkRNA-seq, a Spearman rank correlation analysis was performed between expression values determined by bulkRNA-seq and NanoString (Supplementary Figure 8). Highly significant and strong correlation (rho=0.62-0.88, p-value<2.2e-16) was observed for all sorted cell types (Supplementary File 4). Overall, gene expression estimates in the bulkRNA-seq dataset were confirmed by using the NanoString assay.

### Defining the transcriptomic landscape of porcine PBMCs at single-cell resolution

Single-cells from PBMCs of seven conventional pigs were partitioned, sequenced, clustered, and visualized (Supplementary File 6). In total, the final dataset included 28,810 cells expressing from 9,176-12,683 genes, and each cell was assigned to one of 36 transcriptionally-distinct clusters (Figure 4A, Supplementary Figure 9A-C; Supplementary File 6). For identification of general cell types in each cluster, expression levels of genes known to be active in distinct porcine immune cell populations were mapped across single-cell clusters (Figure 4B-C). The 36 clusters were deduced to 13 general cell types (Figure 4D) as described below.

**Figure 4.**
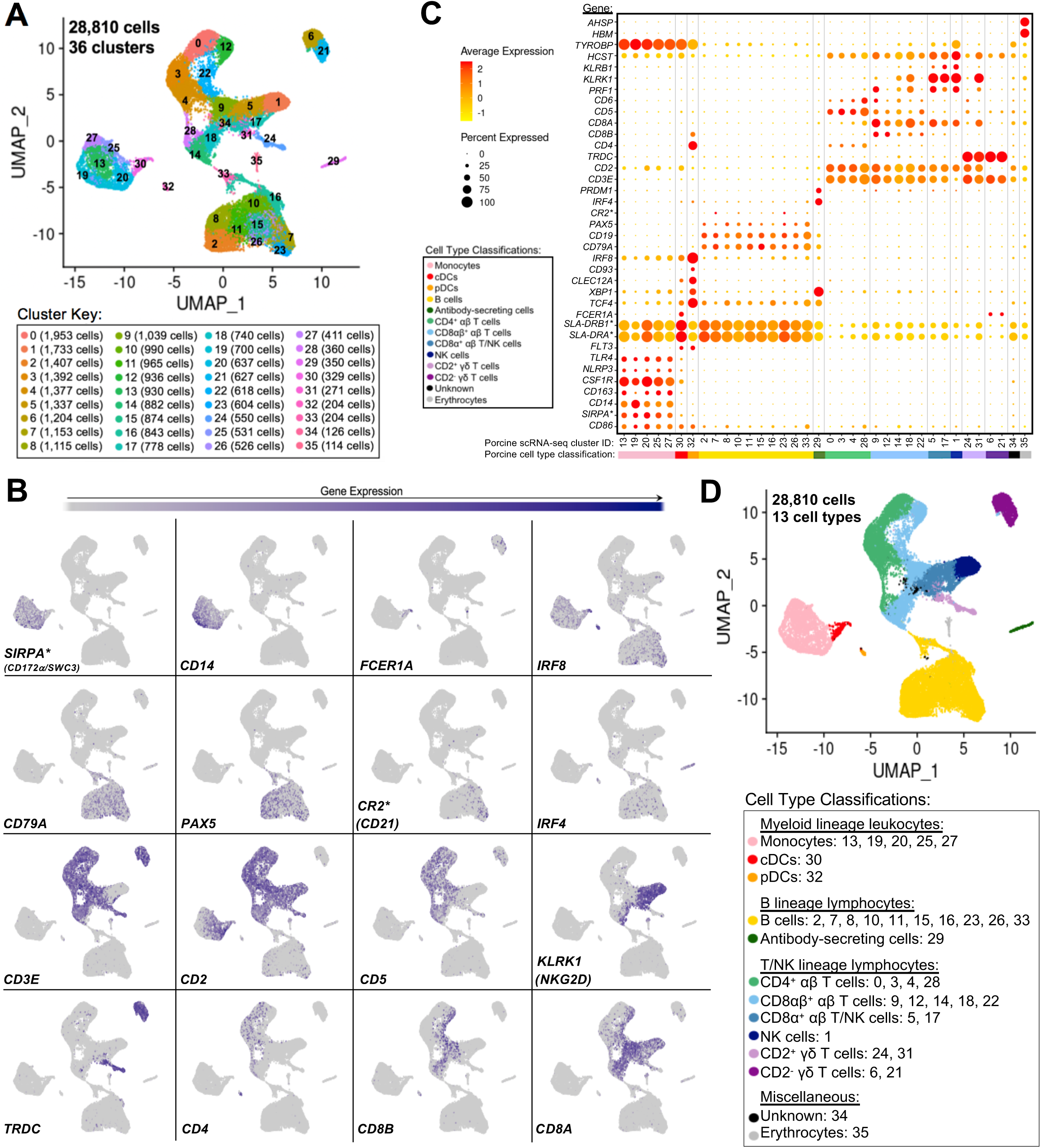
Classification of porcine PBMC scRNA-seq clusters based on known cell type-specific gene expression. **A)** Two-dimensional UMAP visualization of 28,810 single cells from porcine PBMCs classified into 36 designated clusters. Each point represents a single cell. Color of the point corresponds to transcriptional cluster a cell belongs to. Cells more transcriptionally similar to each other belong to the same cluster. **B)** Visualization of selected cell type-specific gene expression overlaid onto two-dimensional UMAP coordinates of single cells. Each point represents a single cell. Color of the point corresponds to relative expression of a specified gene (bottom left of each UMAP plot) within a cell. Grey corresponds to little/no gene expression, while navy corresponds to increased gene expression. **C)** Dot Plot visualization of selected cell type-specific gene expression for each single-cell cluster shown in A. Clusters are listed on the x-axis, while selected genes are listed on the y-axis. The size of a dot corresponds to the percent of cells in a cluster that expressed the gene. The color of a dot corresponds to the average relative expression level for the gene in the cells expressing the gene within a cluster. Color bar below the x-axis corresponds to porcine cell type each cluster was classified as. **D)** Two-dimensional UMAP visualization of single cells from porcine PBMCs classified into major porcine cell types. Each point represents a single cell. Color of the cell corresponds to porcine cell type the respective cluster was designated as based on gene expression patterns for the cluster it belonged to in C. Seven PBMC samples used for scRNA-seq analysis were derived from each of three separate experiments (experiment B, n=2; experiment C, n = 3; experiment D, n = 2). Between 3,042 and 6,518 cells were derived from each PBMC sample. *Refer to ‘Gene name replacement’ methods.

Monocyte clusters (13, 19, 20, 25, 27) expressed *CSF1R* and genes associated with microbial recognition (*CD14, CD163, NLRP3, TLR4*), reported as highly expressed by porcine monocytes (Auray et al., 2016). DC clusters (30, 32) expressed porcine pan-DC marker *FLT3* and were further classified as conventional DCs (cDCs; cluster 30) by elevated expression of *FCER1A* and MHCII-encoding genes (*SLA-DRB1*, SLA-DRA\**) and pDCs (cluster 32) by elevated expression of *TCF4, XBP1, CLEC12A, CD93, IRF8, CD4,* and *CD8B* (Auray et al., 2016). Co-stimulatory gene *CD86* was expressed by all monocyte and DC clusters as reported (Auray et al., 2016). *SIRPA*^*^, encoding CD172α is expressed by porcine monocytes/DCs (Piriou-Guzylack and Salmon, 2008; Auray et al., 2016) and used to sort myeloid leukocytes for bulkRNA-seq above, was minimally expressed in DC clusters.

B-cell clusters (2, 7, 8, 10, 11, 15, 16, 23, 26, 33) expressed *CD79A, CD19,* and *PAX5* (Faldyna et al., 2007; Piriou-Guzylack and Salmon, 2008; Bordet et al., 2019). Antibody-secreting cells (ASCs; cluster 29) expressed *IRF4* and *PRDM,* genes ascribed to immunoglobulin secretion (Shi et al., 2015; Liu et al., 2020). Detection of *CR2\**, the gene encoding CD21 protein, was very low in any cluster.

Expression of *CD3E*, which encodes pan-T-cell CD3ε protein, identified T-cell clusters (0, 3, 4, 5, 6, 9, 12, 14, 17, 18, 21, 22, 24, 28, 31) (Gerner et al., 2009a). Cluster 1 cells largely lacked *CD3E, CD5,* and *CD6* expression, while expressing *CD2, CD8A*, *PRF1,* NK receptor-encoding genes *KLRB1* (CD161) and *KLRK1* (NKG2D), and NK receptor signaling adaptor molecules *HCST* (DAP10) and *TYROBP* (DAP12), corresponding to a NK cell designation (Denyer et al., 2006; Piriou-Guzylack and Salmon, 2008; Gerner et al., 2009a; Toka et al., 2009). γδ T-cells were identified by *TRDC* expression, encoding the γδTCR δ chain, and were subdivided into two major subtypes based on presence/absence of *CD2* expression (Stepanova and Sinkora, 2013; Sedlak et al., 2014) (Piriou-Guzylack and Salmon, 2008; Gerner et al., 2009a). Clusters 6 and 21 were identified as CD2^-^ γδ T-cells and clusters 24 and 31 as CD2^+^ γδ T-cells. Clusters expressing *CD3E* but not *TRDC* were considered αβ T-cells and were further subdivided based on *CD4* expression (0, 3, 4, 28 classified as CD4^+^ αβ T-cells) or *CD8A* and *CD8B* expression (9, 12, 14, 18, 22 classified as CD8αβ^+^ αβ T-cells) (Piriou-Guzylack and Salmon, 2008; Gerner et al., 2009a). Clusters 5 and 17 were more difficult to fully classify and likely represented a mixture of cells, with some but not all cells expressing *CD3E*. Cells in clusters 5 and 17 largely lacked expression of *CD5*, *CD6, TRDC, CD4,* and *CD8B* but did largely express *CD2, CD8A*, *KLRB1*, and *KLRK1* and were therefore characterized as a mixture of CD8α^+^ αβ T- and NK cells.

Cells in cluster 34 could not be characterized well enough to broadly classify as myeloid, B, T, or NK lineage leukocytes based on the porcine cell markers described and remained unclassified. Cluster 35 expressed *HBM* and *AHSP*, indicating erythrocytes. Clusters 34 and 35 were still included in further scRNA-seq analyses; however, results pertaining to these clusters were not discussed.

### Gene signatures of bulkRNA-seq populations had limitations in resolving single-cell identities

Gene set enrichment analyses (GSEA) using SEG lists defined at different levels of enrichment for each sorted bulkRNA-seq population (Supplementary File 3, see Methods) was performed to identify scRNA-seq clusters were likely represented (Figure 5A-B, Supplementary Figure 10A, Supplementary File 8). Some gene sets had high relative enrichment in anticipated corresponding scRNA-seq clusters, such as Myeloid gene sets to monocyte/DC clusters, CD21nB/CD21pB gene sets to B-cell clusters, and SWC6gdT gene sets to CD2^-^ γδ T-cell clusters. Interestingly, highest relative enrichment (2.51) for the top 1% of CD21nB SEGs was noted for ASCs in cluster 29, followed by erythrocytes in cluster 35 (1.68). Within sorted NK and T-cell populations, some gene sets showed high relative enrichment for their anticipated corresponding clusters in the scRNA-seq dataset. We also noted off-target relative enrichment for gene sets in clusters not anticipated to be included in specific sorted cell populations. Cluster 28 had lower relative enrichment for CD4T and CD4CD8T SEG lists at top 5-25% SEG levels (−0.02 to 0.73) than did several non-CD4^+^ αβ T-cell clusters. Similar phenomena were observed for CD8T top 5-25% SEG lists, whereby clusters 1, 24, and 31 had higher relative enrichment for CD8T SEG lists (0.69 to 1.56) than did clusters 14 or 18 (−0.04 to 0.95 relative enrichment) that were anticipated to be included in the CD8T population. Clusters 24 and/or 31 showed off-target relative enrichment for all T/NK gene sets to various degrees, though these cells would not be expected to make up a sizeable portion of any of those sorted cell populations.

**Figure 5.**
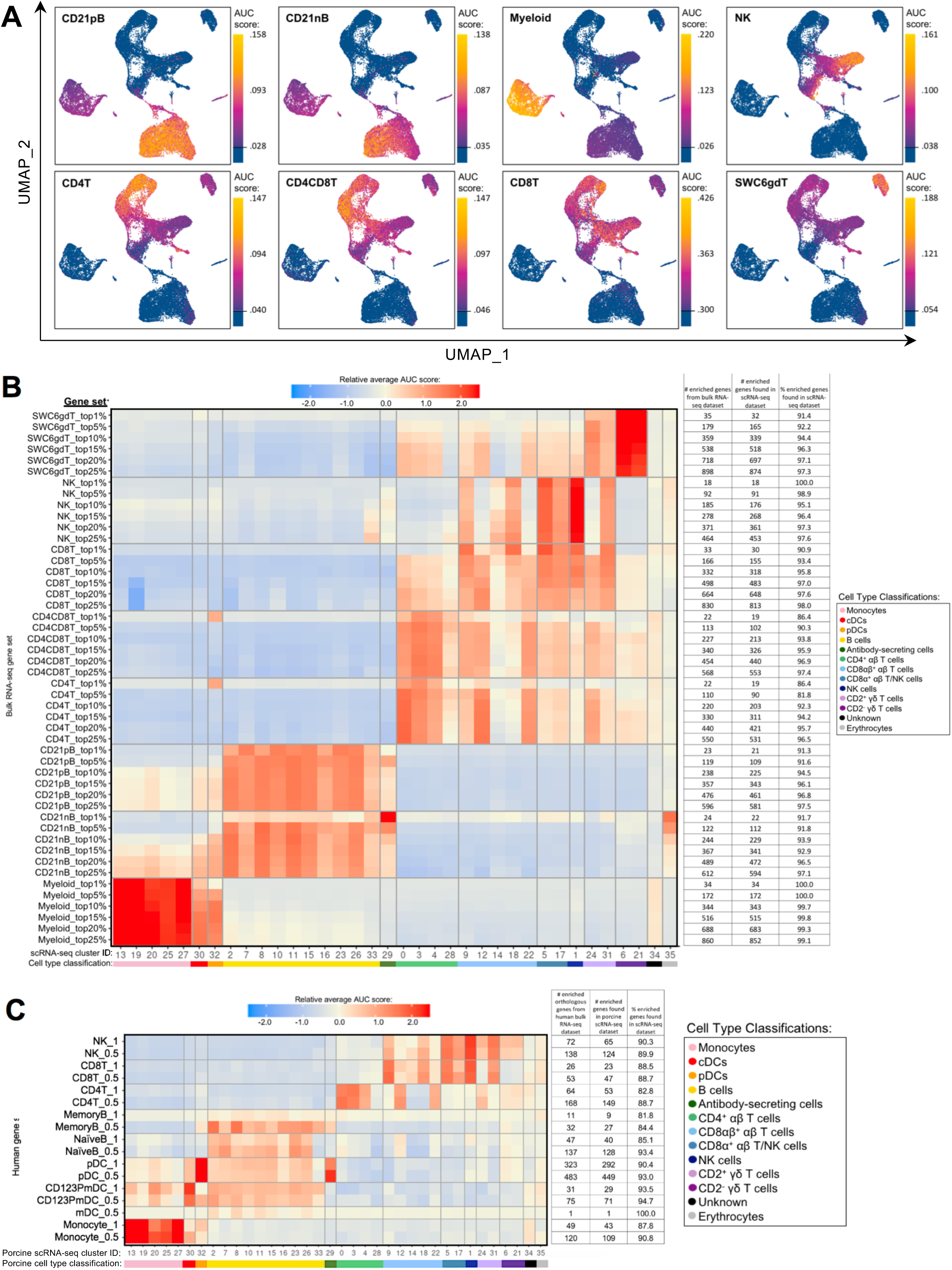
Enrichment of gene signatures from bulkRNA-seq in porcine single-cell clusters. **A)** Gene set enrichment scores calculated by AUCell analysis of enriched gene sets from the top 25% of SEGs in pig bulkRNA-seq sorted populations overlaid onto cells of the porcine scRNA-seq dataset visualized in two-dimensional UMAP plot. Each point represents a single cell. The color of the point corresponds to the AUC score calculated for each respective cell. Higher AUC scores correspond to a greater percentage of cells from a gene set being detected in the top 5% of expressed genes in a cell. A threshold for AUC score detection within each gene set was set as shown in Supplementary Figure 10A and is indicated by a horizontal line on the gradient fill scale for each plot. **B)** Relative average gene set enrichment scores of scRNA-seq clusters calculated by AUCell analysis of enriched gene sets from porcine bulkRNA-seq sorted data. Scores are relative to other cells within a single gene set comparison (across a row of the heatmap) and are not calculated relative to scores across different gene sets (across columns in the heatmap). Gene sets were created from the top 1, 5, 10, 15, 20, or 25% of SEGs from sorted populations, as determined by highest log2FC values in the porcine bulkRNA-seq data. The number of genes included from the bulkRNA-seq dataset and the number and percent of genes detected in the scRNA-seq dataset is listed on the right of the heatmap. A color bar under scRNA-seq cluster IDs indicates the cell type classification, as according to Figure 4D. **C)** Relative average gene set enrichment scores of scRNA-seq clusters calculated by AUCell analysis of enriched gene sets from human bulkRNA-seq sorted data. Scores are relative to other cells within a single gene set comparison (across a row of the heatmap) and are not calculated relative to scores across different gene sets (across columns in the heatmap). Gene sets were created from genes with high expression scores > 0.5 or >1 for each respective sorted population of cells, with a greater high expression score indicating greater enrichment. The number of genes included from the bulkRNA-seq dataset and the number and percent of genes detected in the scRNA-seq dataset is listed on the right of the heatmap. A color bar under scRNA-seq cluster IDs indicates the cell type classification, as according to Figure 4D.

Further comparison of porcine bulk and scRNA-seq data by CIBERSORTx deconvolution analysis largely supported our single-cell cluster designations by predicting which clusters proportionally represented the bulk RNA-seq data. (Supplemental Figure 10 B, Supplemental File 7). Several clusters with poor AUCell enrichment for anticipated bulkRNA-seq gene sets in Figure 5A-B, such as cluster 28, were predicted to constitute considerable proportions of their anticipated cell populations by CIBERSORTx deconvolution analysis. Additionally, clusters that demonstrated off-target enrichment by AUCell analysis, such as clusters 1, 9, 22, 24, and 31, were not predicted to be largely present in those off-target populations using CIBERSORTx. However, CIBERSORTx failed to predict many single-cell clusters to have notable abundances in any bulkRNA-seq populations, such as clusters 8, 19, 26, 32, and 34 having < 3.33% predicted abundance for any one bulkRNA-seq sample.

Additional GSEA comparing gene sets derived from public bulkRNA-seq data of sorted human PBMC populations with porcine single-cell gene expression profiles informed cluster identity as it relates to human immune cells (Figure 5C, Supplementary Figure 10C-D, Supplementary File 9). High relative enrichment for human monocyte gene sets in porcine monocyte populations, human CD123PmDC gene sets in porcine cDCs, and human pDC gene sets in porcine pDCs was observed, in general consensus with gene expression profiles of anticipated corresponding porcine single-cell clusters. NaiveB gene signatures had positive relative enrichment in all porcine B-cell clusters except cluster 33 at both the 0.5 and 1.0 resolution level, while the MemoryB signature had highest relative enrichment scores for B and ASC clusters at the 0.5 level, with little relative enrichment at the 1.0 level (likely due to a limited number of genes in the gene set). Human T/NK gene sets had off-target enrichment very similar to patterns observed in GSEA with porcine gene sets. Overall, GSEA between human bulkRNA-seq gene signatures and gene expression profiles of porcine scRNA-seq data supported many of the same findings when comparing between porcine bulkRNA-seq gene sets and gene expression profiles of porcine scRNA-seq data. Results indicated limitations of gene profiles obtained from sorted bulkRNA-seq populations in accurately describing/accounting for transcriptional heterogeneity resolved by scRNA-seq.

### Integration of porcine and human scRNA-seq datasets to further annotate porcine cells

We examined porcine single-cell identities by comparing the porcine scRNA-seq data to a highly annotated scRNA-seq dataset of human PBMCs, providing a higher level of resolution than available with bulkRNA-seq. Transfer of more highly-specified human cell type labels onto porcine cells could reveal the most likely human counterparts for these porcine populations. Mapping scores were further calculated to determine how well porcine cells were truly represented by the human dataset (Figure 6A, Supplementary Figure 11A-B, Supplementary File 10.

**Figure 6.**
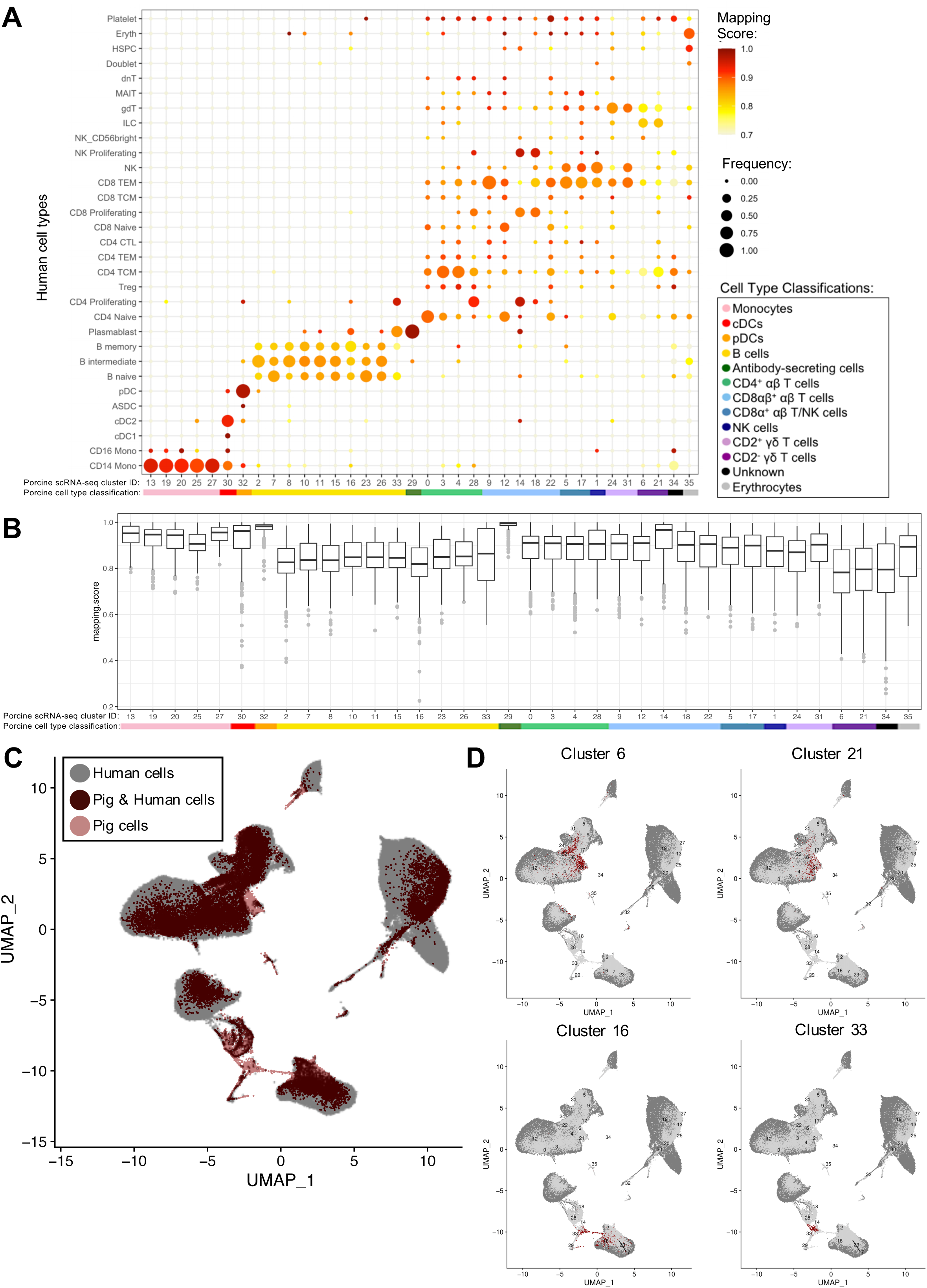
Integration of porcine and human scRNA-seq datasets to further annotate porcine cells**. A)** Mapping scores calculated to determine how well porcine cells were represented by the human dataset. The human cell type specific frequency (size of the circle) and mapping score for that human cell type (color) are shown for each porcine scRNA-seq cluster. Porcine cell type classifications (color) are shown below the porcine scRNA-seq cluster IDs. **B)** Mapping scores calculated to determine how well porcine cells were represented by the human dataset. The mapping scores for each porcine scRNA-seq cluster is represented by a box and whiskers plot. Porcine cell type classifications (color) are shown below the porcine scRNA-seq cluster IDs. **C)** To identify cells in the porcine dataset that were not well represented in the human dataset, a de-novo visualization of the merged porcine and human data was performed. The porcine (pink) and human (grey) were plotted together using UMAP. An overlap of both porcine and human cells is shown as (dark red). Clusters of porcine cells that are not well represented in the human data can be observed by pink regions in the plot. **D)** Two primary regions of porcine cells that were not well represented in the human data were identified in C. In order to clarify which porcine scRNA-seq clusters were represented in these regions, the porcine cluster IDs were projected onto the UMAP and cells from four clusters overlapping the identified regions were colored as dark red.

Many porcine clusters had >95% of cells mapping to a specific human cell type, with average mapping scores >0.9, including monocyte, pDC, cDC, and ASC clusters, suggesting high congruency between pig and human for these cell types (Figure 6B). All porcine B-cell clusters, omitting cluster 33, mapped primarily to human B-cell clusters, but average mapping scores were slightly lower (0.80-0.87), indicating less ideal representation in the human data. In addition, every porcine B-cell cluster had overlap with all three human B-cell types (Figure 6A). Of the porcine CD4^+^ αβ T-cells, most cluster 0 cells were predicted as human CD4 naïve cells, clusters 3 and 4 cells as human CD4 T central memory (TCM) cells, and cluster 28 cells as human CD4 proliferating cells. From porcine CD8αβ^+^ αβ T-cells, clusters 14 and 18 were largely assigned as human replicating cell types, while 90% of cluster 9 cells were predicted as human CD8 T effector memory (TEM) cells. Highest cluster 12 predictions were mainly to human CD4/CD8 naïve T-cells, and cluster 22 cells predicted to match a range of human cell populations, with the largest percentage predicted as human CD8 TEMs. Porcine CD8α^+^ αβ T/NK and NK clusters had predictions split primarily across human CD8 TEM and NK designations. Porcine CD2^+^ γδ T-cell clusters 24 and 31 had 74% and 98%, respectively, of cells predicted as human CD8 TEM, NK, or γδ T-cells. Porcine CD2^-^ γδ T-cell clusters 6 and 21 had the majority of cells predicted as human CD4 TCM, innate lymphoid cell (ILC), or γδ T-cells, though the average mapping scores were lower for those assigned as CD4 TCM (0.73-0.74) or gdT (0.74-0.78) than those assigned as ILCs (0.82-0.83) (Supplementary File 10). Overall, cross-species comparison to a well-annotated human scRNA-seq dataset helped elucidate porcine cell type identities at a higher resolution than porcine or human bulkRNA-seq datasets (Figure 5), though some discordance was clearly still present.

Several porcine clusters had low mapping scores to a human cell type, indicating the porcine cells may not be well represented by the human reference dataset (Figure 6B and Supplementary File 10). Therefore, *de novo* visualization was performed on the combined human and porcine data, to identify cells in the pig dataset not well represented in the human data (Figure 6C-D). Porcine clusters could be identified that had low similarity to human cells, and vice versa (Figure 6C). Specifically, porcine clusters 6, 16, 21, and 33 weakly overlapped human cells in the two-dimensional *de novo* visualization (compare 6C and 6D). Furthermore, clusters 6, 16, 21, and 33 had lower average mapping scores to any human cell type (Figure 6B).

### Different activation states of porcine CD4^+^ αβ T-cells based on CD8α expression

We further compared scRNA-seq gene expression profiles amongst only CD4^+^ αβ T-cell clusters to gain functional inferences and correspondence to CD8α^-^ versus CD8α^+^ phenotypes that were used to sort CD4^+^ αβ T-cells for bulkRNA-seq. CD4^+^ αβ T-cell clusters (0, 3, 4, 28) were comprised of 5,082 total cells (Figure 7A). Hierarchical clustering and pairwise DGE (Supplementary File 7), as well as random forest (RF) analyses, a deep-learning classification method, (see Methods; Supplementary File 11), cumulatively revealed clusters 3 and 4 to be the most transcriptionally similar to each other. Clusters 3 and 4 had the smallest hierarchical distance, fewest DEGs (67), and largest RF error rate (19.5) between them, while cluster 28 was the most distantly related to the other 3 clusters (Figure 7B).

**Figure 7.**
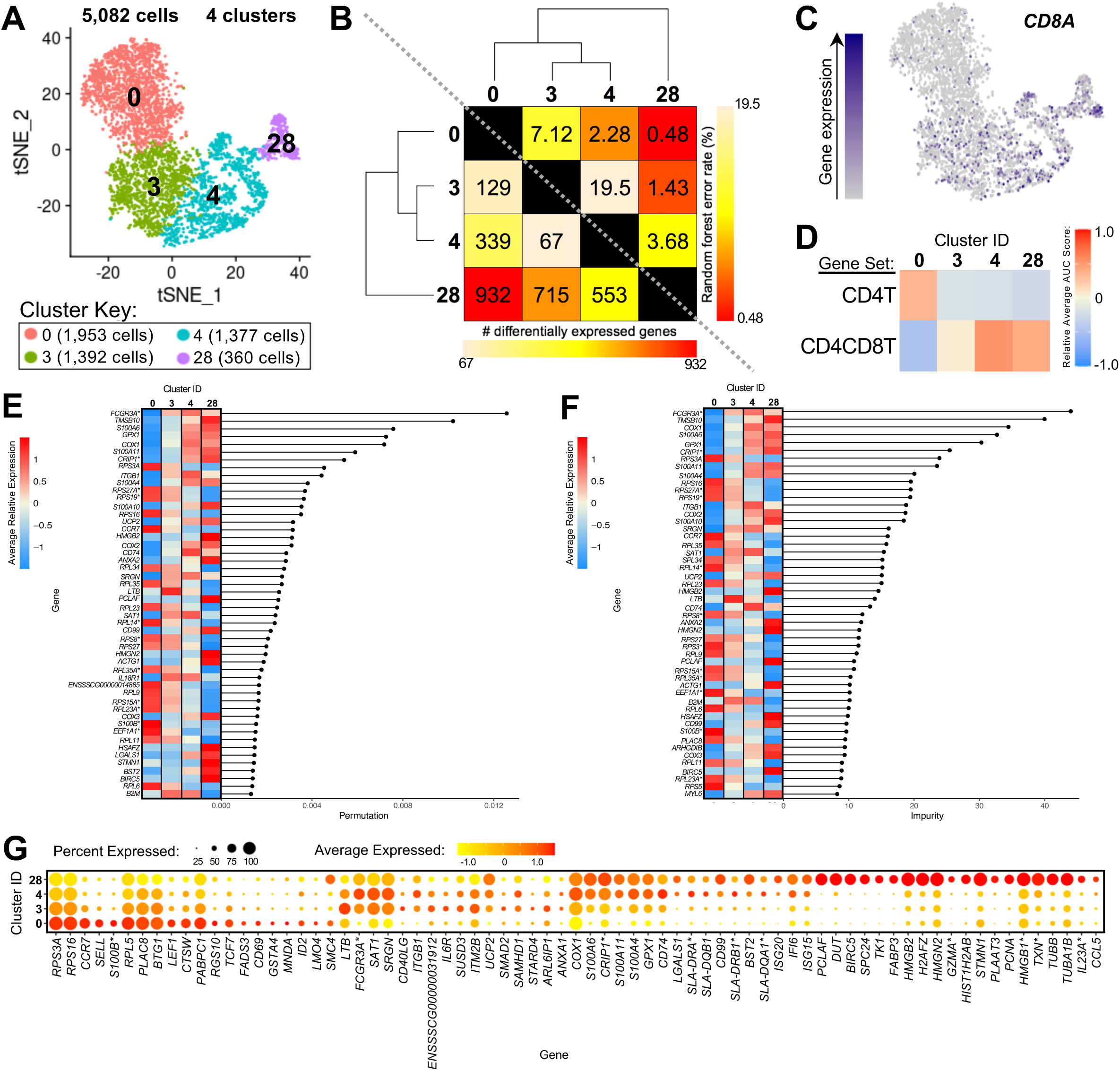
Transcriptional heterogeneity of porcine CD4+ ab T-cells at single-cell resolution. **A)** Two-dimensional t-SNE plot of 5,082 cells belonging to clusters designated as CD4+ ab T-cells (clusters 0, 3, 4, 28) in Figure 4D. Each point represents a single cell. Color of the cell corresponds to transcriptional cluster a cell belongs to. Cells more transcriptionally similar to each other belong to the same cluster. **B)** Transcriptomic relationship amongst CD4+ ab T-cell clusters as calculated by three methods: hierarchical clustering (as seen by hierarchical trees on both axes), pairwise random forest analyses (as seen on top right diagonal); and pairwise DGE analyses (as seen on bottom left diagonal). Longer branches on the hierarchical tree corresponds to greater hierarchical distance. Lower numbers of DEGs by DGE analysis and higher out-of-bag (OOB) error rates from random forest analyses indicate greater pairwise transcriptional similarity. **C)** Visualization of CD8A expression overlaid onto t-SNE coordinates of single CD4+ ab T-cells. Each point represents a single cell. Color of the point corresponds to relative expression of CD8A within a cell. Grey corresponds to little/no gene expression, while navy corresponds to increased gene expression. **D)** Relative average gene set enrichment scores of CD4+ ab T-cell clusters calculated by AUCell analysis of DEG sets from pairwise DGE analysis of the CD4T and CD4CD8T populations from porcine bulkRNA-seq. Scores are relative to other cells within a single gene set comparison (across a row of the heatmap) and are not calculated relative to scores across gene set (across columns in the heatmap). **E&F)** Genes with the largest effects in discriminating CD4+ ab T-cells by cluster identities were determined, as indicated by high permutation **(E)** and/or impurity scores **(F)** calculated from a trained random forest model. Average relative expression for each of these genes within clusters is also depicted by a heatmap. **G)** Dot plot of up to the top 20 DEGs having logFC > 0 from overall DGE analysis of only CD4+ ab T-cell clusters. Clusters are listed on the y-axis, while selected DEGs are listed on the x-axis. The size of a dot corresponds to the percent of cells in a cluster that expressed the gene. The color of a dot corresponds to the average relative expression level for the gene in the cells expressing the gene within a cluster. *Refer to ‘Gene name replacement’ methods.

*CD8A* gene expression was detected in a subset of cells in the CD4^+^ αβ T-cell clusters (3.5%, 13.1%, 20.9%, 39.7% of cells in clusters 0, 3, 4, 28, respectively; Figure 7C). *CD8A* expression was significantly greater in clusters 4 and 28 compared to cluster 0 by pairwise DGE analyses (Supplementary File 7) but not in cluster 3 compared to 0, due to not meeting a minimum threshold of cells (20%) expressing the gene in either cluster implemented for DGE analysis. However, cluster 3 had significantly greater expression of *CD8A* compared to cluster 0 when removing the minimum cell expression threshold (average log2FC =0.37, adjusted p-value =5.52×10-21). GSEA of DEGs identified by pairwise DGE analysis of CD4T and CD4CD8T populations recovered from bulkRNA-seq (Supplementary File 2) revealed genes significantly enriched in CD4T compared to CD4CD8T populations were relatively enriched in cluster 0, while genes significantly enriched in CD4CD8T compared to CD4T populations showed greater relative enrichment in clusters 4 and 28 and to a lesser extent in cluster 3 (Figure 7D, Supplementary File 12).

The top genes contributing to overall transcriptional heterogeneity amongst four clusters of CD4^+^ αβ T-cells, as determined by RF analysis (Figure 7 E-F, Supplementary File 13), highly overlapped with genes identified in overall DGE analysis (Figure 7G, Supplementary File 13). Of eight genes with mutually highest permutation and impurity scores from overall RF analysis (Figure 7E-F), one gene had significantly greater expression in cluster 0 compared to all other clusters (*RPS3A*), while the other seven genes had significantly greater expression in clusters 3, 4, and 28 compared to cluster 0 (*FCGR3A*,TMSB10, COX1, S100A6, GPX1, CRIP1*, S100A11*), as determined by pairwise DGE analyses (Supplementary File 7).

Genes associated with a naïve phenotype, including *CCR7, SELL, LEF1,* and *TCF7* (Szabo et al., 2019; Kim et al., 2020) had significantly increased expression in cluster 0 (Figure 7G, Supplementary File 9 and 13), in line with the result obtained by comparing to human scRNA-seq data that indicated a good alignment of cluster 0 with human naïve CD4 T-cells (Figure 6A). From Figure 6A, clusters 3 and 4 aligned with human CD4 Tcm (central memory) cells, and cluster 28 aligned with human CD4 proliferating cells. Correspondingly, genes associated with activation, such as *ITGB1, CD40LG, IL6R,* and MHC II-associated genes (*CD74, SLA-DRA, SLA-DQB1, SLA-DRB1*, SLA-DQA1*^*^) (Grewal and Flavell, 1996; Gerner et al., 2009b; Zemmour et al., 2018; Zhu et al., 2020) had significantly greater expression in clusters 3, 4, and/or 28, and cluster 28 expressed many genes specific for cellular replication and division (*PCLAF, BIRC5, TK1, PCNA*) (Dabydeen et al., 2019; Giotti et al., 2019) (Figure 7G, Supplementary File 9 and 13). Overall, we leveraged single-cell gene expression profiles to confirm likely identity of cluster 0 as naïve CD4^+^CD8α^-^ αβ T-cells and clusters 3, 4, and 28 as potentially previously activated CD4^+^CD8α^+^ αβ T-cells.^1^

### Heterogeneity between/amongst CD2^+^ and CD2^-^ γδ T-cells

Clusters predicted to be porcine γδ T-cells were examined to reveal transcriptional distinctions within this cell type. Four clusters containing 2,652 cells were previously identified as CD2^-^ γδ T-cells (clusters 6, 21) or CD2^+^ γδ T-cells (clusters 24, 31) (Figure 8A). We could further segregate these clusters by *CD2* and *CD8A* expression into CD2^-^CD8α^-^ (clusters 6, 21), CD2^+^CD8α^-^ (cluster 24), and CD2^+^CD8α^+^ (cluster 31) designations used to functionally define porcine γδ T-cells previously (Stepanova and Sinkora, 2013; Sedlak et al., 2014) (Figure 8B).

**Figure 8.**
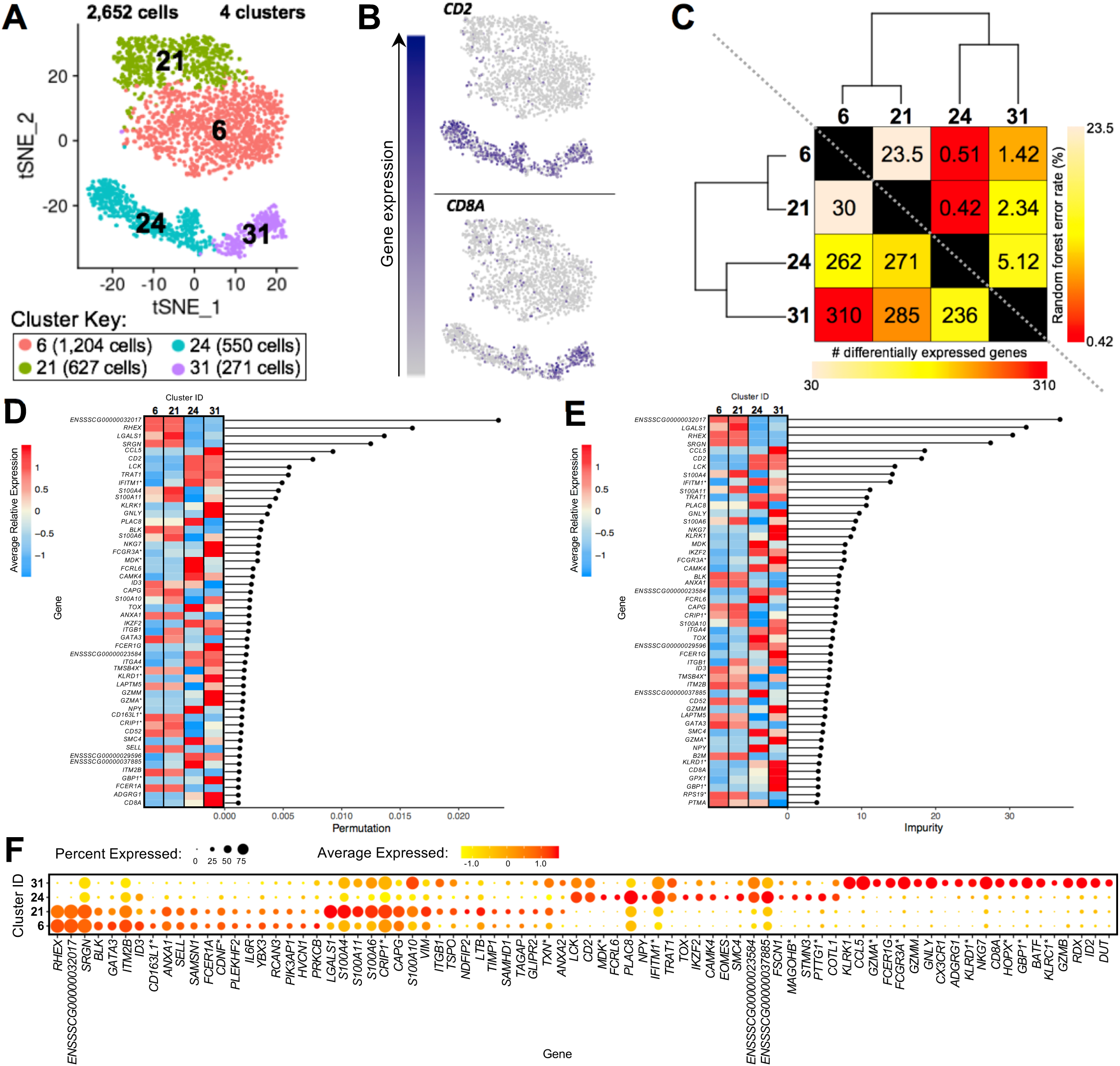
Transcriptional heterogeneity of porcine gd T-cells at single-cell resolution. **A)** Two-dimensional t-SNE plot of 2,652 cells belonging to clusters designated as CD2-gd T-cells (clusters 6, 21) or CD2+ gd T-cells (clusters 24, 31) in Figure 4D. Each point represents a single cell. Color of the cell corresponds to transcriptional cluster a cell belongs to. Cells more transcriptionally similar to each other belong to the same cluster. **B)** Visualization of selected gene expression overlaid onto t-SNE coordinates of single gd T-cells. Each point represents a single cell. Color of the point corresponds to relative expression of a specified gene (top left of each t-SNE plot) within a cell. Grey corresponds to little/no gene expression, while navy corresponds to increased gene expression. **C)** Transcriptomic relationship amongst gd T-cell clusters as calculated by three methods: hierarchical clustering (as seen by hierarchical trees on both axes), pairwise random forest analyses (as seen on top right diagonal); and pairwise DGE analyses (as seen on bottom left diagonal). Longer branches on the hierarchical tree corresponds to greater hierarchical distance. Lower numbers of DEGs by DGE analysis and higher out-of-bag (OOB) error rates from random forest analyses indicate greater pairwise transcriptional similarity. **D&E)** Genes with the largest effects in discriminating gd T-cells by cluster identities were determined, as indicated by high permutation **(D)** and/or impurity scores **(E)** calculated from a trained random forest model. Average relative expression for each of these genes within clusters is also depicted by a heatmap. **F)** Dot plot of up to the top 20 DEGs having logFC > 0 from overall DGE analysis of only gd T-cell clusters. Clusters are listed on the y-axis, while selected DEGs are listed on the x-axis. The size of a dot corresponds to the percent of cells in a cluster that expressed the gene. The color of a dot corresponds to the average relative expression level for the gene in the cells expressing the gene within a cluster. *Refer to ‘Gene name replacement’ methods.

CD2^-^ γδ T-cell clusters 6 and 21 were most closely related to one another by hierarchical clustering, had the fewest pairwise DEGs (30), and had the highest pairwise RF analysis error rate (23.5), indicating clusters 6 and 21 to be the most transcriptionally similar γδ T-cell clusters of the four clusters (Figure 8C, Supplementary File 7 and 14). CD2^+^ γδ T-cell clusters 24 and 31 were most similar to each other by hierarchical clustering, had the second fewest pairwise DEGs (236), and had the second highest pairwise RF error rate (5.12), indicating clusters 24 and 31 to be most similar to each other. When performing pairwise comparison between any CD2^-^ and CD2^+^ γδ T-cell clusters, the number of DEGs increased and RF error rates decreased, indicating greater transcriptional differences between cells of the CD2^-^ and CD2^+^ γδ T-cell lineages than amongst them (Figure 8C, Supplementary File 7 and 14).

The top genes contributing to overall transcriptional heterogeneity amongst γδ T-cell clusters, as determined by RF analysis (Figure 8D-E, Supplementary File 14), overlapped with genes identified with significant and highest logFC expression in overall DGE analysis (Figure 8F, Supplementary File 14). Six of the top seven genes with mutual highest impurity (the best features that correctly split the data) and permutation scores from RF analysis (Figure 8D-E) were also DEGs between both CD2^-^ compared to both CD2^+^ γδ T-cell clusters by pairwise DGE analysis (Supplementary File 7), again indicating large transcriptional differences between CD2^-^ and CD2^+^ γδ T-cells. In total, 31 genes had significantly greater expression in both CD2^-^ γδ T cell clusters compared to both CD2^+^ γδ T cell clusters, and 49 genes had significantly greater expression in both CD2^+^ γδ T cell clusters compared to both CD2^-^ γδ T cell clusters (Table 4), as determined using the pairwise DGE analyses (Supplementary File 7).

**Table 4.**
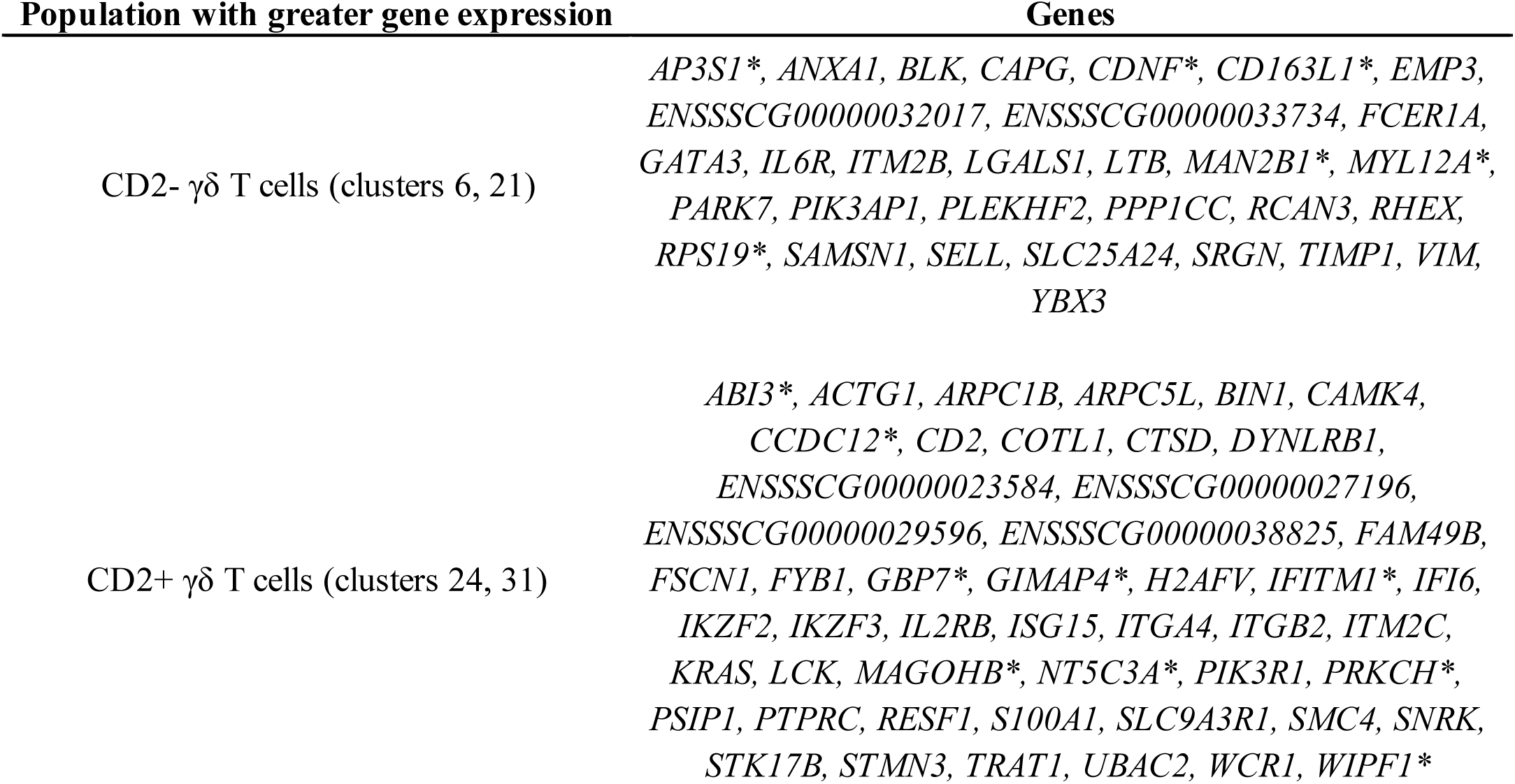
Genes differentially expressed between both CD2-gd T-cell clusters (clusters 6, 21) and both CD2+ gd T-cell clusters (clusters 24, 31). * Refer to gene name replacement in methods

Intra-lineage heterogeneity of CD2^-^ γδ T-cells (between clusters 6 and 21) and CD2^+^ γδ T-cells (between clusters 24 and 31) demonstrated additional complexity beyond the inter-lineage heterogeneity between CD2^-^ and CD2^+^ γδ T-cells. Pairwise comparison between clusters 24 and 31 (Supplementary Data 8) revealed 80 genes with significantly greater expression in cluster 24 (CD2^+^CD8α^-^ γδ T-cells) and 156 genes with significantly greater expression in cluster 31 (CD2^+^CD8α^+^ γδ T-cells). Genes with the greatest logFC expression (logFC > 1.5) in cluster 31 compared to cluster 24 were related to cellular activation and/or effector functions (*CCL5, GNLY, FCGR3A*, KLRK1, GZMA*, NKG7, FCER1G, GZMB*) (Rincon-Orozco et al., 2005; Pizzolato et al., 2019; Szabo et al., 2019). Of the 30 DEGs between clusters 6 and 21 (Supplementary Data 8), three genes had significantly greater expression in cluster 6, while 27 genes had significantly greater expression in cluster 21. Several genes with greater expression in cluster 21 encoded for activation- or stress-induced molecules, including *GPX1, LGALS1, ITGB1, LTB,* several genes encoding for S100 proteins (*S100A4, S100A6, S100A10, S100A11*), and genes related to MHCII presentation (*CD74, SLA-DRA*^*^) (Blaser et al., 1998; Ware, 2005; Gerner et al., 2009b; Steiner et al., 2011; Kesarwani et al., 2013; Siegers, 2018). Genes encoding transcriptional regulators playing important roles in cell fate determination, including *ID3* and *GATA3,* had greater expression in cluster 6, while *ID2* expression was significantly greater in cluster 21 (Blom et al., 1999; Zhang et al., 2014; Rodríguez-Gómez et al., 2019).^1^

## DISCUSSION

We present the first comprehensive annotation of the global transcriptome of all major circulating porcine blood mononuclear cells. We applied bulkRNA-seq to determine transcriptomes of eight sorted PBMC populations and scRNA-seq to annotate transcriptomic diversity of PBMCs into transcriptionally-distinct clusters. Deep RNA sequencing detected significant heterogeneity between sorted populations except for T cell populations, while further heterogeneity was unmasked by scRNA-seq. Collectively, the data sets revealed specific immune functional expression patterns and highlighted substantial diversity in some subsets, such as T-cells. The combined approach helps to unite porcine transcriptomics and cellular immunology, as transcriptional differences and functional relationships of porcine immune cells have remained unclear due to lack of sufficient reagents to label distinct porcine immune cell populations. While cross-species comparisons have been done with many RNA-seq datasets of partially purified cell populations (Kapetanovic et al., 2013; Herrera-Uribe et al., 2020), our new porcine data demonstrates global similarity to human bulkRNA-seq and scRNA-seq transcriptomes that can be used to further unravel porcine cell function and extend comparative immune investigation.

Gene expression patterns from the bulkRNA-seq datasets revealed distinct transcript profiles enriched in biological pathways characteristic of each respective cell population, based on previous findings in pig and other species (Alter et al., 2004; Palmer et al., 2006; Wang et al., 2008; Foissac et al., 2019; Monaco et al., 2019; Summers et al., 2020). However, bulkRNA-seq data from the porcine sorted populations had limited ability to identify genes with specific transcriptional patterns for some sorted lymphocyte populations. The transcriptomes of eight different cell types we provide include three types of transcriptomes that have not reported before in pig, including NK, CD21pB and CD21nB. Lists of SEGs, pairwise DGE between all populations and cell type-specific genes data sets presented here, could be used for further analysis in other pig or even in cross-species comparisons. Notably, we were able to identify a large number of HEGs in the Myeloid population. Some HEGs in Myeloid cells were reported as a Myeloid cell markers in pig (e.g. *CD14* and *CD36*) (Fairbairn et al., 2013) and other HEGs may be considered as new potential cell markers. Also, in comparison to sorted CD4T and CD8T populations reported in a previous porcine RNA-seq study (Foissac et al., 2019), we observed concordant transcriptional patterns in essentially equivalent populations. However, we extended transcriptional annotation to two additional T-cell populations (CD4CD8T, SWC6gdT), thus identifying transcriptional differences across more T-cell populations. We demonstrated the utility of an established NanoString CodeSet (Van Goor et al., 2020; Dong et al., 2021) to validate RNA-seq results and further profile porcine sorted PBMC populations. At the bulk RNAseq level, we concluded substantial transcriptional heterogeneity was present across sorted T-cell and B-cell populations, as fewer enriched or cell type-specific genes were detected. As described below, the lack of identification of cell type-specific genes was likely caused by the lack of further sub-setting during sorting to separate functionally distinct cells. However, we were able to find several specific transcriptional patterns in B- and T-cells using bulkRNA-seq, and some of the identified genes encode for transmembrane proteins. Beyond further description of well-annotated genes, we also demonstrated that up to 18% of our predicted cell-type specific and enriched genes are currently poorly annotated, i.e., genes with no recognized human ortholog. These data thus increase the functional annotation of these genes, as co-expression patterns linking such genes with known genes can be an important component for Gene Ontology classifications and disease-association gene prediction (van Dam et al., 2018), and is an important proposed outcome of the FAANG project (Giuffra et al., 2019).

Comparison of our sorted population expression patterns to a similar human RNA-seq dataset revealed both similarities and differences between species. While we compared the transcriptomes of the sorted cells with human populations that were isolated using similar cell markers, we cannot exclude that we are biasing this comparison due to different immunoreagent markers used across species. However, we did find similar transcriptional patterns across immune cell populations that are intrinsic to a lineage, such as the porcine Myeloid population correlating with the human myDC123 population, in agreement with other studies (Auray et al., 2016).

Previous global gene expression studies using either porcine whole blood or specific immune cell types have failed to thoroughly describe all major PBMC populations (Freeman et al., 2012; Dawson et al., 2013; Mach et al., 2013; Auray et al., 2016; Foissac et al., 2019). Providing the transcriptomes of bulk sorted cell populations will be readily useful to the majority of porcine immunology research labs that use sorting techniques to analyze porcine immune cell function and RNA expression patterns, as new lists of co-expressed genes in these cell populations are now available. However, our combined analysis of such bulkRNAseq data with the scRNAseq data demonstrated that the former approach has significant heterogeneity, limiting the ability to resolve specific cell types for deeper transcriptional interrogation. A combined analysis provided evidence confirming our hypothesis that scRNA-seq would lead to identification of more specific and novel transcriptional signatures to improve annotation and understanding of circulating porcine immune cells.

scRNA-seq provides many noted benefits in transcriptomic analysis, however there are limitations to the approach. Of benefit, scRNA-seq captured transcriptomes of cells excluded from our bulkRNA-seq analysis, as scRNA-seq approach did not rely on protein marker expression and selection of sorting criteria based on specific marker phenotypes. As mentioned above, scRNA-seq also established that greater levels of cellular heterogeneity exist, since sequencing was resolved to the level of individual cells rather than a sorted population. We recognize the scRNAseq-predicted clusters may contain transitory cell states that may be very challenging to further study for the relationship between cellular function and transcriptional patterns (Bassler et al., 2019). Further, we assumed single-cell gene expression profiles would be indicative of protein expression for cell type-specific markers; however, gene expression for many such markers, including *SIRPA\** and *CR2\** that encode proteins used for bulk RNA-seq cell sorting, was sparse. Sparsity of data is a known limitation of the scRNA-seq approach utilized herein, while methods such as imputation have been proposed to improve sensitivity (Andrews et al., 2021). We chose not to use imputation due to our current inability to estimate effects on cell patterns through comparison to an external reference (Andrews et al., 2021). Thus, these limitations made it difficult to decipher between low- and non-expression for some genes of interest, including canonical markers used for identifying cell types in the immunology literature. Instead, reliance on gene expression profiles of multiple markers was used. For example, *SIRPA*^*^ expression was observed at low levels in monocyte clusters but was virtually absent in DC clusters, though both porcine monocytes and DCs express CD172A protein. Because DCs express CD172A at lower levels than monocytes (Piriou-Guzylack and Salmon, 2008; Auray et al., 2016), *SIRPA\** expression in DCs may have been below our limit of detection using scRNA-seq, as it was insufficiently expressed in DCs but not in monocytes. We utilized a droplet-based partitioning method for scRNA-seq that can detect a large number of cells but a lower number of transcripts per cell. By this method, we could retain a large number of cells (>25,000 cells from seven samples) at the expense of limited sequencing depth per cell (minimum of 500 unique genes and 1,000 unique transcripts per cell). Utilizing higher sequencing depth per cell or different partitioning platforms for scRNA-seq that have more efficient transcript capture per cell will be beneficial for deeper analysis of specific cells/genes of interest. It is likely some gene expression profiles are not predictive of protein expression, due to post-transcriptional regulation mechanisms. Using newly available co-expression lists to formulate more refined cell sorting regimens and scRNAseq analysis of such sorted populations will also increase the ability to define transcriptomes of such cell types (Nestorowa et al., 2016). It was notable that the lists of genes predicted to be significantly enriched in the 36 scRNAseq clusters had overall a very similar fraction of poorly annotated genes (average of 18%; cite in Supplementary File 6) to those predicted for bulkRNAseq, indicating that even the genes with expression patterns predicted to be more discriminatory contribute a similar level of genome annotation improvement.

We used multiple methods to compare these high-dimensional expression datasets to further interpret genes predicted to be different between sorted cell populations, between clusters, or between human and pig. GSEA and/or deconvolution analyses of bulkRNA-seq to scRNA-seq datasets was only partially effective in correlating sorted populations with assumed corresponding clusters in the scRNA-seq dataset (regardless of inter-species or intra-species comparison). At a higher level of resolution, both methods were able to assign most corresponding cell-type designations between scRNA-seq and bulkRNA-seq data. However, several different scRNA-seq clusters were not predicted to make up a large portion of any bulkRNA-seq sample. While methodology could account for these differences, it is more likely that CIBERSORTx was unable to discriminate between certain clusters due to their high similarity. For example, cells that could have been predicted to be assigned to cluster 8, which makes up a large proportion of the scRNA-seq data, may have been assigned to other similar B cell clusters. The ability to discriminate between similar clusters may have been impacted by down sampling each cluster to include the same number of cells for the analysis. Overall, deconvolution was useful in assigning cell type level data but in some instances, it could not fully deconvolute bulk RNAseq to the cluster specific level.

Integration of porcine PBMC scRNA-seq with a human PBMC scRNA-seq dataset did allow further resolution of porcine cluster annotations and yielded high confidence of homology between many porcine and human single cell populations. While we cannot completely discount the potential for recognized cell types in our scRNA-seq dataset are not present in sorted populations used for bulkRNA-seq (or vice-versa), it seems more likely this is similar evidence to that described above indicating that the same level of resolution simply was not captured by bulkRNA-seq and could not well represent all cell types found in the scRNA-seq data. Integration with another scRNA-seq dataset, even when accounting for cross-species comparison, was in many ways more informative for further annotating porcine single cells, highlighting the enhanced ability of scRNA-seq to define cellular landscapes. Moreover, cross-species integration extended our knowledge of comparative immunology between humans and pigs, as we could identify most similar human counterparts by reference-based prediction. Conversely, we could also identify clusters of CD2^-^ γδ T-cells (clusters 6 and 21) and B-cells (clusters 16 and 33) that were largely specific to the porcine dataset by *de novo* visualization of clustered cells using the combined human and pig data. The clusters of cells in pig samples either represent porcine cells either lacking close human cellular counterparts or the equivalent human counterparts were not well-captured in the human PBMC scRNA-seq dataset.

While we did not perform deeper biological query of all cell types identified in our scRNA-seq dataset, we did attempt to deduce biological significance for the different CD4^+^ αβ T-cell populations that have unique aspects in pigs. Deeper query of CD4^+^ αβ T-cells was performed, as there is functional interest in determining activation states of porcine CD4^+^ αβ T^-^ cells based on CD8α expression, which may be gained upon activation and retained in a memory state (Summerfield et al., 1996; Zuckermann, 1999; Saalmüller et al., 2002; Gerner et al., 2009b). We found it difficult to identify CD4^+^ αβ T-cell clusters as CD8α^+^ or CD8α^-^ due to sparsity in *CD8A* expression but could leverage comparison of CD4T and CD4CD8T populations from bulkRNA-seq to formulate gene sets enriched in each CD4 expressing T-cell population. GSEA helped identify one cluster of CD4^+^CD8α^-^ αβ T-cells that corresponded mostly to human naïve CD4 T-cells, while three clusters of CD4^+^CD8α^+^ αβ T-cells corresponded to human memory or proliferating CD4 T-cells. Collectively, these data reinforce previous porcine literature, elucidate parallels to human cells, and provide greater insight into the spectrum of activation states present in CD4^+^CD8α^+^ αβ T-cells. Future analysis of activated T-cells or trajectory analysis may provide even further insight on the transition of activation states in porcine peripheral T-cells.

Pigs are a ‘γδ high’ species, named as such because they have a higher proportions of γδ T-cells in circulation, largely attributed to the presence of CD2^-^ γδ T-cells that are absent in humans and mice (Stepanova and Sinkora, 2013). Three major γδ T-cell populations are characterized in pigs: CD2^-^CD8α^-^ γδ T-cells that express SWC6 and CD2^+^CD8α^-/+^ γδ T-cells that do not express SWC6, where CD2^-^CD8α^-^ γδ T-cells become CD2^+^CD8α^+^ upon activation (Stepanova and Sinkora, 2013; Sedlak et al., 2014). As our sorting strategy for bulkRNA-seq utilized an anti-SWC6 antibody rather than a pan-γδ T-cell-specific antibody; thus, γδ T-cells for bulk RNA-seq included CD2^-^CD8α^-^ γδ T-cells in the SWC6gdT population or CD2^+^CD8α^+^ γδ T-cells found in combination with CD4^-^CD8α^+^ αβ T-cells in the CD8T population. CD2^+^CD8α^-^ γδ T-cells were expected to be excluded in cell sorting. In future sorting strategies, it may be beneficial to utilize a pan-γδ T-cell reactive antibody and/or identify CD4^-^CD8^+^ αβ T-cells with anti-CD8β antibody, which should not label with CD2^+^CD8α^+^ γδ T-cells (Gerner et al., 2009b). though this may still exclude potential CD4^-^CD8α^+^CD8β^-^ αβ T cells, such as we observed in clusters 5 and 17. Despite limitations in sorting, the bulkRNA-seq profiles were still informative when comparing to scRNA-seq data. The highest relative enrichment of SWC6gdT gene signatures was detected in CD2^-^ γδ T-cell clusters, while CD2^+^ γδ T-cell clusters showed relative enrichment to a lesser level, indicating some conserved gene expression between CD2^-^and CD2^+^ γδ T-cells. Comparison between CD2^+^ γδ T-cell clusters further supported previous biological understanding, where CD2^+^CD8α^+^ γδ T-cells had greater expression of genes related to cellular activation and cytotoxicity relative to CD2^+^CD8α^-^ γδ T-cells (Yang and Parkhouse, 1997; Stepanova and Sinkora, 2013; Sedlak et al., 2014). On the other hand, CD2^-^ γδ T-cells are less well described than CD2^+^ γδ T-cells, largely due to lack of comparable populations in humans or mice that may be used for biological inference. Integration with human scRNA-seq data supported previous observations of the absence of CD2^-^ γδ T-cells in humans, as close counterparts for CD2^-^ γδ T-cell clusters could not be found by *de novo* visualization, and reference-based integration indicated closest human counterparts to be a mixture of primarily γδ T-cells, ILCs, and CD4 T_CM_s, and mapping scores were highest for human ILCs rather than γδ T-cells, indicating human ILCs to be the closest, albeit still poor, human match. Nonetheless, we were able to highlight transcriptional distinctions that better annotate CD2^-^ γδ T-cells, including DEGs between CD2^-^ and CD2^+^ γδ T-cells that defined the two γδ T cell lineages and between two clusters of CD2^-^ γδ T-cells that have not yet been described.

## Conclusion

This study provides a first-generation atlas annotating circulating porcine immune cell transcriptomes at both the cell surface marker-sorted population and single-cell levels. These findings illuminate the landscape of immune cell molecular signatures useful for porcine immunology and a deeper annotation of the genome, a goal of the FAANG project. These results also provide useful resources to identify new porcine cell biomarkers for discrimination and isolation of specific cell types, urgently needed in the field.

## Supporting information

SuppFile12

SuppFile13

SuppFile10

SuppFile09

SuppFile11

SuppFile14

SuppFile08

SuppFile02

SuppFile05

SuppFile06

SuppFile01

SuppFile04

SuppFile07

SuppFile03

## Abbreviations

AUC: area under the curve
ASC: antibody-secreting cell
B: B-cell
bulkRNA-seq: bulk RNA sequencing
cDC: conventional dendritic cell
DC: dendritic cell
DEGs: differentially expressed genes
DGE: differential gene expression
Exp: experiment
FAANG: Functional Annotation of Animal Genomes
FACS: Fluorescent activated cell sorting
G2P: Genome-to-Phenome
GO: gene ontology
GSEA: gene set enrichment analysis/analyses
HBSS: Hank’s balanced salt solution
HEGs: highly enriched genes
MACS: Magnetic activated cell sorting
mDC/myDC: myeloid dendritic cell
n: negative
NK: natural killer
p: positive
PBMC: peripheral blood mononuclear cell
PC: principal component
PCA: principal component analysis
pDC: plasmacytoid dendritic cell
RF: random forest
RIN: RNA integrity number
RNA-seq: RNA sequencing
scRNA-seq: single-cell RNA sequencing
scREF-matrix: single-cell reference matrix
SEG: significantly enriched genes
sPCA: supervised principal component analysis
SWC6: swine workshop cluster 6
T: T-cell
TCR: T-cell receptor
TPM: transcripts per million
t-SNE: t-distributed stochastic neighbor embedding
UMAP: uniform manifold approximation and projection
UMI: unique molecular identifier
γδ: Gamma-delta
αβ: alpha beta

## Acknowledgments

The authors acknowledge the DNA facility of the Iowa State University for provision of technical support and sequencing platforms utilized in this study. We are grateful to the NADC animal care staff for their efforts. We thank Dr. Catherine Ernst at Michigan State University for providing the Yorkshire pigs used in the bulkRNA-seq study. We thank Samuel Humphrey for cell sorting technical assistance. We thank Kristen Walker for performing the NanoString data collection and initial analyses. We thank Zahra Bond for technical assistance with cell isolation and FACS, and Dr. Julian Trachsel for early data visualization assistance.

## Author Contributions Statements

JH, JEW, KAB, and CLL collected samples and isolated cells. KAB performed cell staining and FACS. JH performed bulk RNA isolations. JL supervised the NanoString assay data collection. JH, LD, and HL performed bulkRNA-seq analyses. HL performed NanoString analyses. JEW, LD, SKS, and HL performed scRNA-seq analyses. JH, JEW, CLL, and CKT interpreted the data and drafted the manuscript. All authors contributed to the writing of the materials and methods, edited the manuscript, and approved the final version.

## Data Availability Statement

Raw sequencing data from bulkRNA-seq and scRNA-seq are available through the European Nucleotide Archive (project: PRJEB43826), https://www.ebi.ac.uk/ena/browser/view/PRJEB43826.

## Footnotes

1 https://www.haemosphere.org/datasets/show

1 http://www.bioinformatics.babraham.ac.uk/projects/fastqc

2 https://jgi.doe.gov/data-and-tools/bbtools/bb-tools-user-guide/bbmap-guide/

1 https://www.haemosphere.org/searches

1 https://cran.r-project.org/web/packages/caret/caret.pdf

2 https://cran.r-project.org/web/packages/ranger/index.html

* Refer to gene name replacement in Methods section

1 http://biocc.hrbmu.edu.cn/CellMarker/#

* Refer to gene name replacement in Methods section

* Refer to gene name replacement in methods

* Refer to gene name replacement in methods

* Refer to gene name replacement in Methods section

