## Supplementary material for "Reference transcriptomes of porcine peripheral immune cells created through bulk and single-cell RNA sequencing": SupplFigs

### Supplementary figures

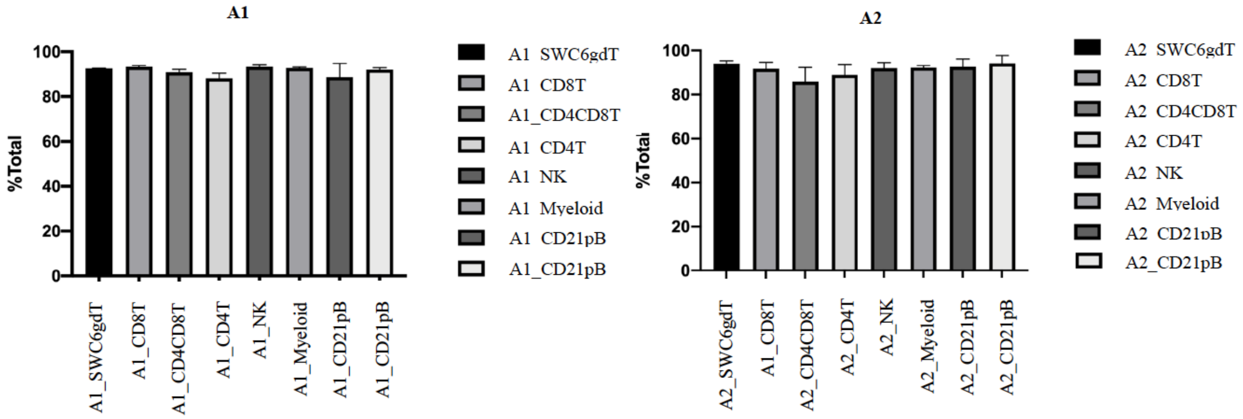


**Supplementary Figure 1.** Percentages for purity yield of sorted cells from two porcine males using BD FACSymphony™.

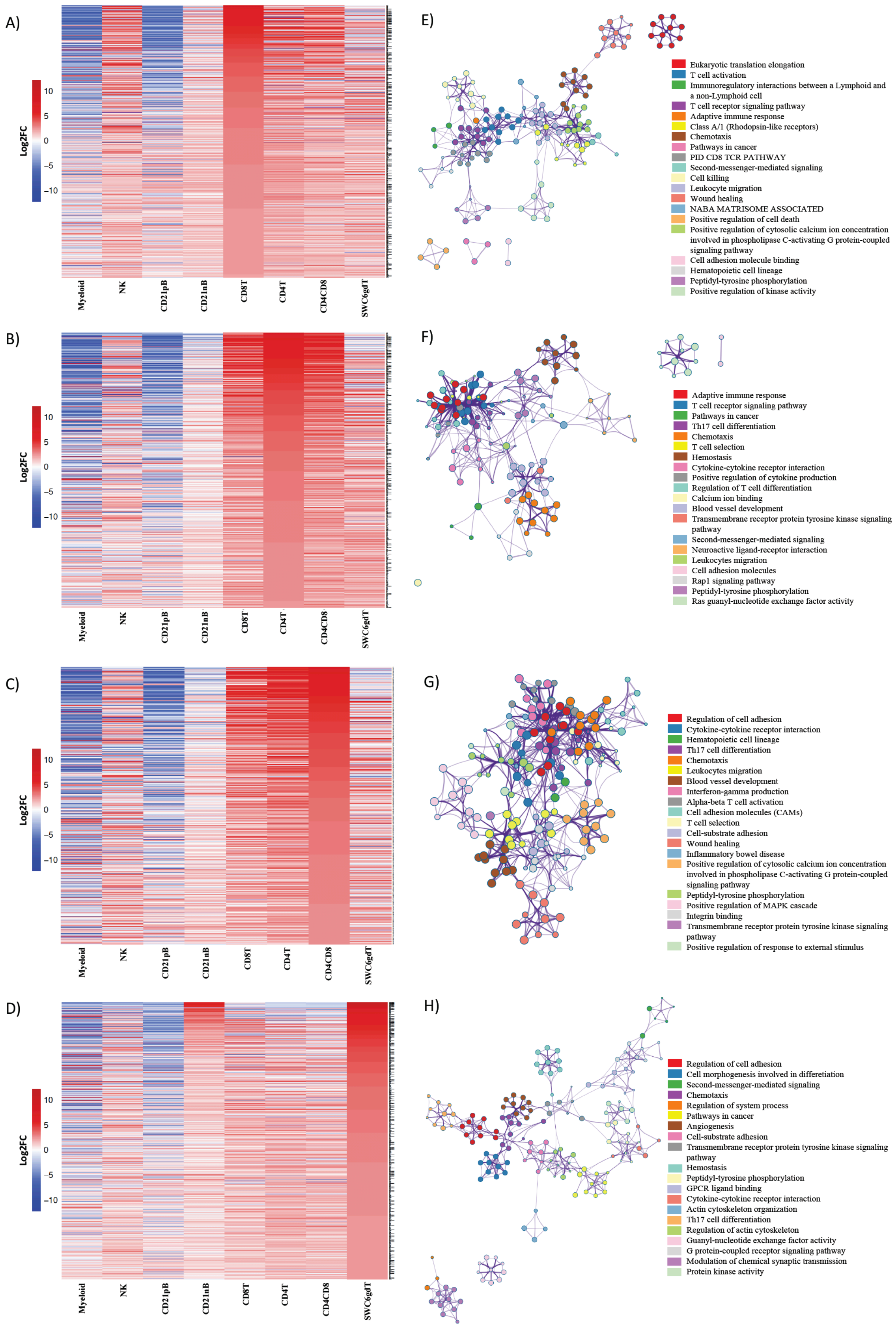
A)
B)
C)
D)
E)
F)
G)
H)
Log2FC
10
5
0
−5
−10
Myeloid
NK
CD21pB
CD21nB
CD8T
CD4T
CD4CD8
SWC6gdT
Eukaryotic translation elongation
T cell activation
Immunoregulatory interactions between a Lymphoid and a non-Lymphoid cell
T cell receptor signaling pathway
Adaptive immune response
Class A/1 (Rhodopsin-like receptors)
Chemotaxis
Pathways in cancer
PID CD8 TCR PATHWAY
Second-messenger-mediated signaling
Cell killing
Leukocyte migration
Wound healing
NABA MATRISOME ASSOCIATED
Positive regulation of cell death
Positive regulation of cytosolic calcium ion concentration involved in phospholipase C-activating G protein-coupled signaling pathway
Cell adhesion molecule binding
Hematopoietic cell lineage
Peptidyl-tyrosine phosphorylation
Positive regulation of kinase activity
Adaptive immune response
T cell receptor signaling pathway
Pathways in cancer
Th17 cell differentiation
Chemotaxis
T cell selection
Hemostasis
Cytokine-cytokine receptor interaction
Positive regulation of cytokine production
Regulation of T cell differentiation
Calcium ion binding
Blood vessel development
Transmembrane receptor protein tyrosine kinase signaling pathway
Second-messenger-mediated signaling
Neuroactive ligand-receptor interaction
Leukocytes migration
Cell adhesion molecules
Rap1 signaling pathway
Peptidyl-tyrosine phosphorylation
Ras guanyl-nucleotide exchange factor activity
Regulation of cell adhesion
Cytokine-cytokine receptor interaction
Hematopoietic cell lineage
Th17 cell differentiation
Chemotaxis
Leukocytes migration
Blood vessel development
Interferon-gamma production
Alpha-beta T cell activation
Cell adhesion molecules (CAMs)
T cell selection
Cell-substrate adhesion
Wound healing
Inflammatory bowel disease
Positive regulation of cytosolic calcium ion concentration involved in phospholipase C-activating G protein-coupled signaling pathway
Peptidyl-tyrosine phosphorylation
Positive regulation of MAPK cascade
Integrin binding
Transmembrane receptor protein tyrosine kinase signaling pathway
Positive regulation of response to external stimulus
Regulation of cell adhesion
Cell morphogenesis involved in differetiation
Second-messenger-mediated signaling
Chemotaxis
Regulation of system process
Pathways in cancer
Angiogenesis
Cell-substrate adhesion
Transmembrane receptor protein tyrosine kinase signaling pathway
Hemostasis
Peptidyl-tyrosine phosphorylation
GPCR ligand binding
Cytokine-cytokine receptor interaction
Actin cytoskeleton organization
Th17 cell differentiation
Regulation of actin cytoskeleton
Guanyl-nucleotide exchange factor activity
G protein-coupled receptor signaling pathway
Modulation of chemical synaptic transmission
Protein kinase activity

**Supplementary Figure 2.** Top 25% highly enriched genes in CD3$^+$ sorted cells. Heatmap showing in decreasing order the top 25% of highly enriched genes in **A**) CD8T, **B**) CD4T, **C**) CD4CD8T and **D**) SWC6gdT cells. Ontology enrichment clusters of the top 25% highly enriched genes of **E**) CD8T, **F**) CD4T, **G**) CD4CD8T and **H**) SWC6gdT cells. The most statistically significant term within similar term clusters was chosen to represent the cluster. Term color is given by cluster ID and the size of the terms is given by –log10 P-value. The stronger the similarity among terms, the thicker the edges between them.

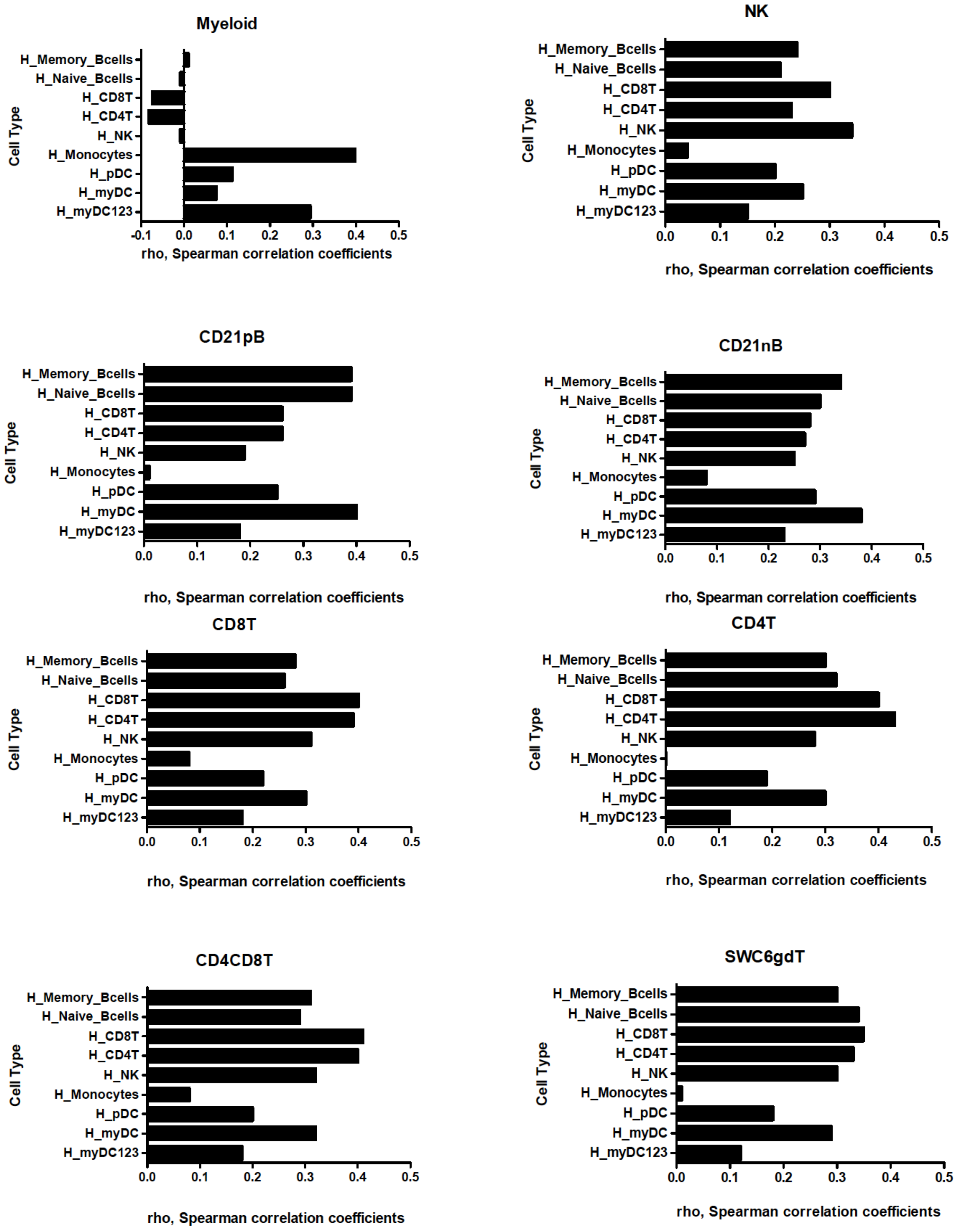


**Supplementary Figure 3.** Spearman rank correlation between TPM values of expressed genes in porcine sorted immune cells and their orthologous genes in human sorted immune cells.

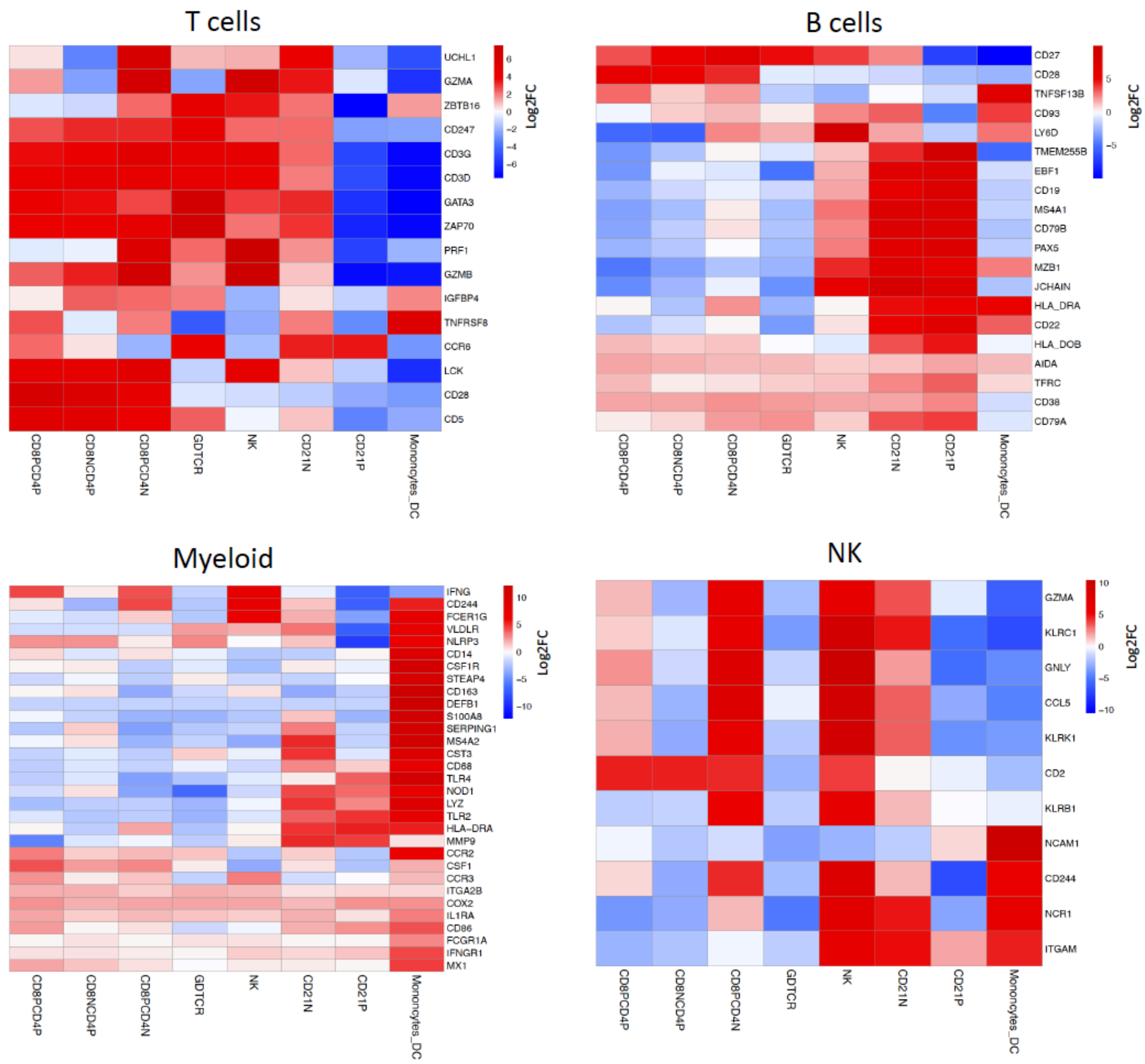


**Supplementary Figure 4.** Gene enrichment heatmap of highly expressed genes and cell markers reported in human and mouse databases in T cells, B cells, myeloid and NK cells.

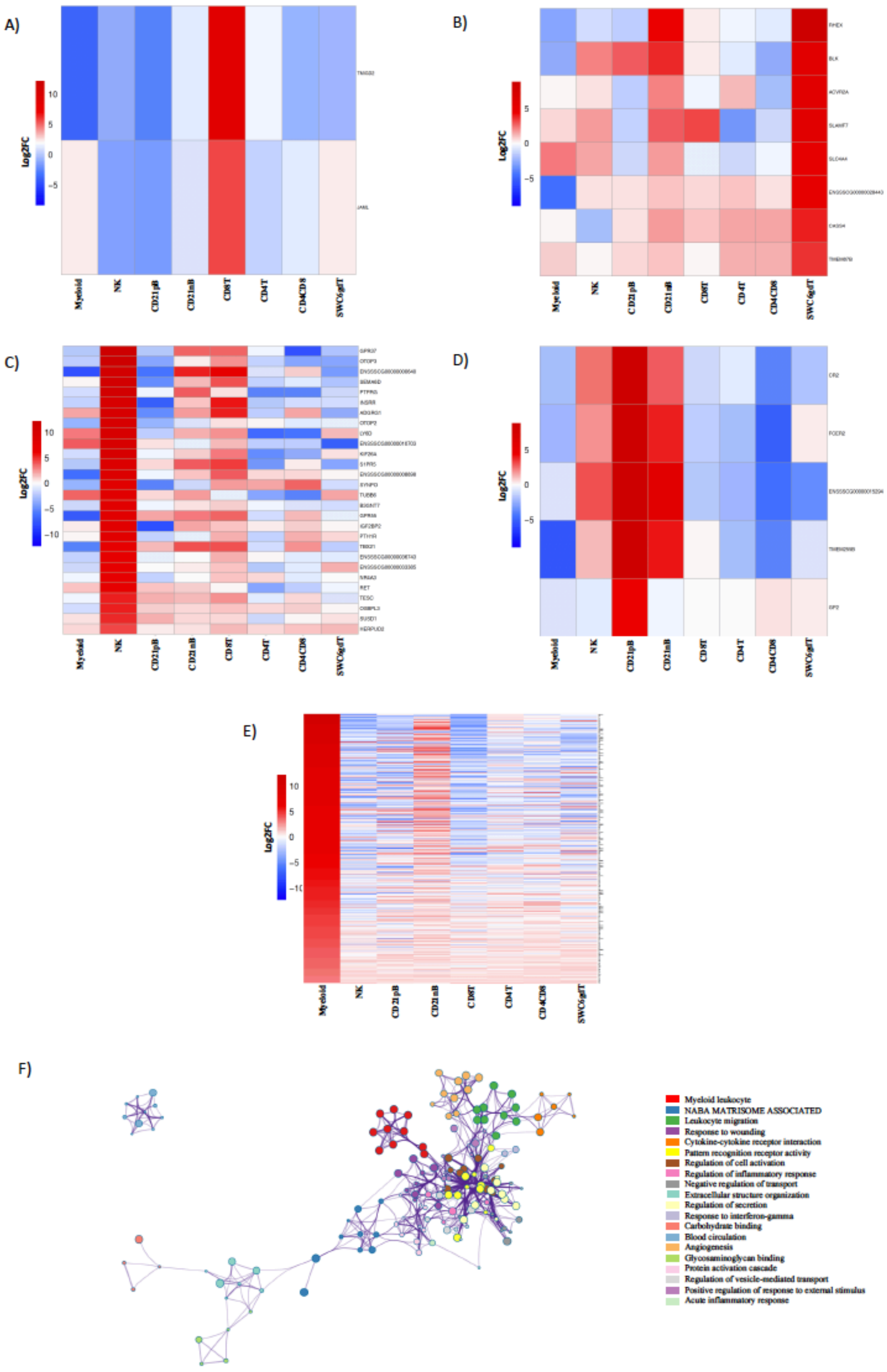

A)
B)
C)
D)
E)
F)
Log2FC
Myeloid
NK
CD21pB
CD21nB
CD8T
CD4T
CD4CD8
SWC6gdT
Myeloid leukocyte
NABA MATRISOME ASSOCIATED
Leukocyte migration
Response to wounding
Cytokine-cytokine receptor interaction
Pattern recognition receptor activity
Regulation of cell activation
Regulation of inflammatory response
Negative regulation of transport
Extracellular structure organization
Regulation of secretion
Response to interferon-gamma
Carbohydrate binding
Blood circulation
Angiogenesis
Glycosaminoglycan binding
Protein activation cascade
Regulation of vesicle-mediated transport
Positive regulation of response to external stimulus
Acute inflammatory response

**Supplementary Figure 5.** Cell-type specific genes in CD8T, SWC6gdT, NK, CD21pB and myeloid cells. Heatmap showing in decreasing order the cell-type specific genes in **A**) CD8T, **B**) SWC6gdT, **C**) NK, **D)** CD21pB and **E)** myeloid. Ontology enrichment clusters cell-type specific genes of **F**) myeloid cells. The most statistically significant term within similar term cluster was chosen to represent the cluster. The most statistically significant term within similar term clusters was chosen to represent the cluster. Term color is given by cluster ID and the size of the terms is given by –log10 P-value. The stronger the similarity among terms, the thicker the edges between them.

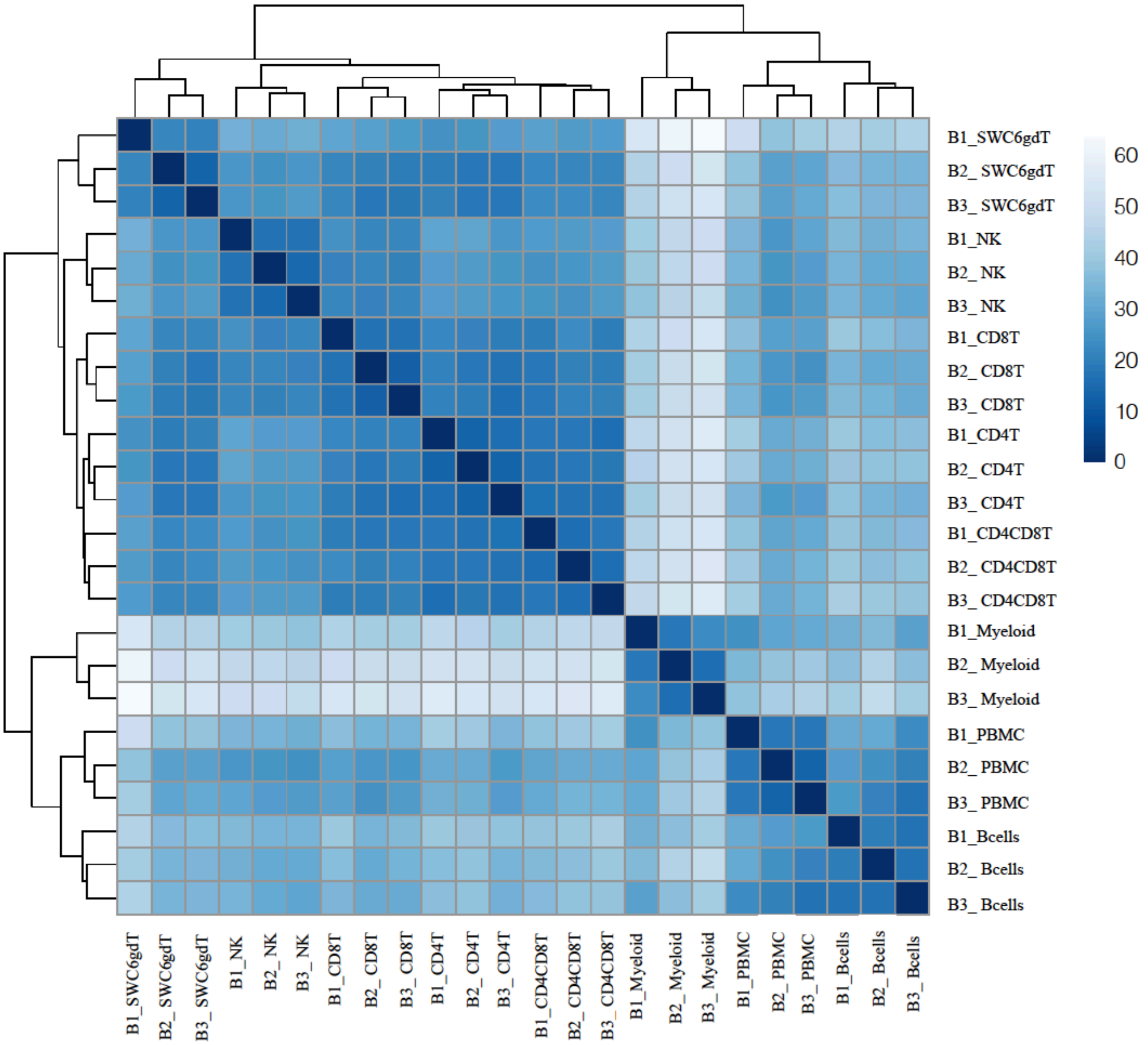


**Supplementary Figure 6.** Gene expression patterns of sorted immune cells showed closer clustering than those share far common progenitors during hematopoiesis. Principal component analysis of transformed reads counts for selected genes in Nanostring. Axis indicate component scores.

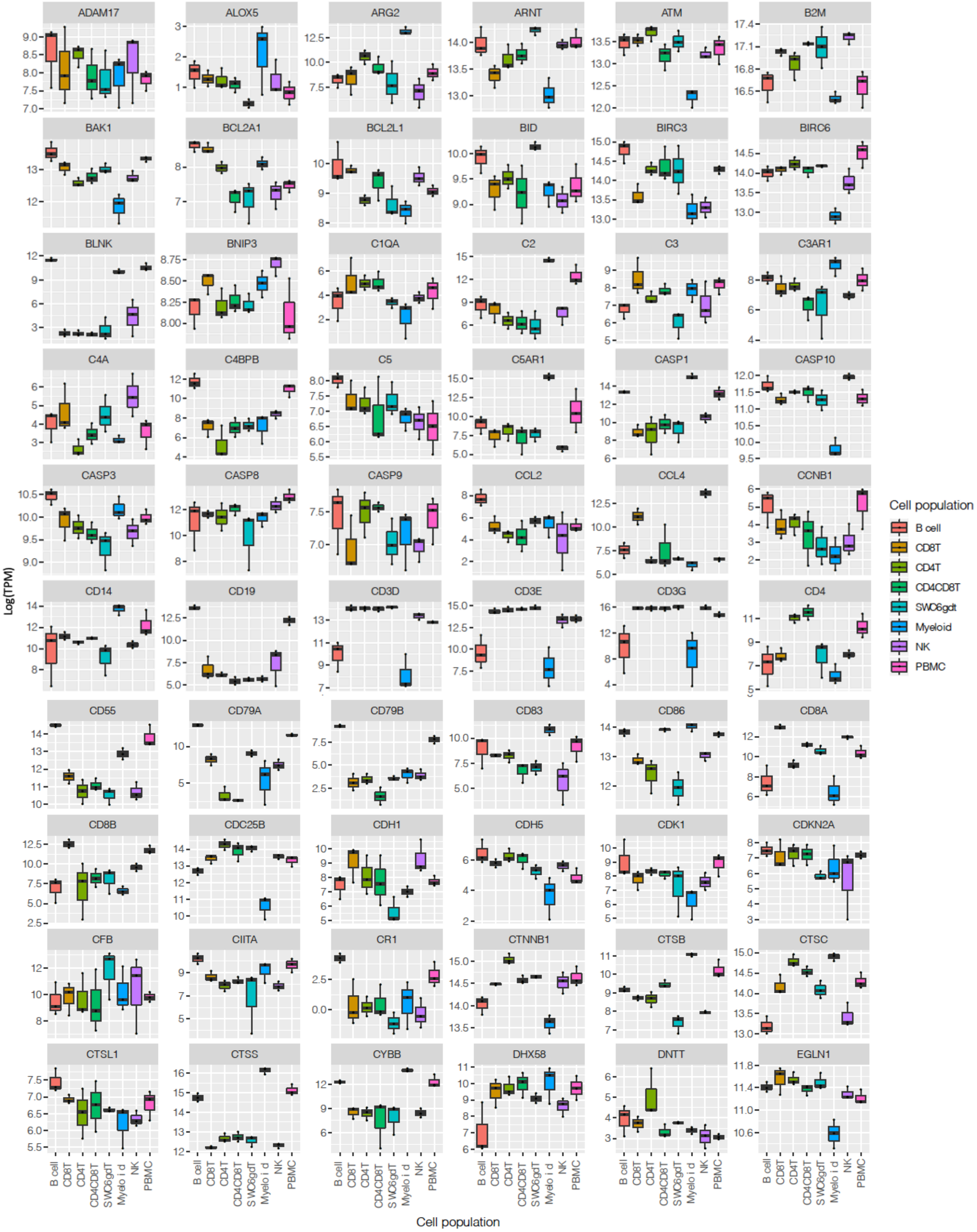
ADAM17
ALOX5
ARG2
ARNT
ATM
B2M
BAK1
BCL2A1
BCL2L1
BID
BIRC3
BIRC6
BLNK
BNIP3
C1QA
C2
C3
C3AR1
C4A
C4BPB
C5
C5AR1
CASP1
CASP10
CASP3
CASP8
CASP9
CCL2
CCL4
CCNB1
CD14
CD19
CD3D
CD3E
CD3G
CD4
CD55
CD79A
CD79B
CD83
CD86
CD8A
CD8B
CDC25B
CDH1
CDH5
CDK1
CDKN2A
CFB
CIITA
CR1
CTNNB1
CTSB
CTSC
CTSL1
CTSS
CYBB
DHX58
DNTT
EGLN1
Log(TPM)
Cell population
B cell
CD8T
CD4T
CD4CD8T
SWC6gdt
Myeloid
NK
PBMC
Cell population

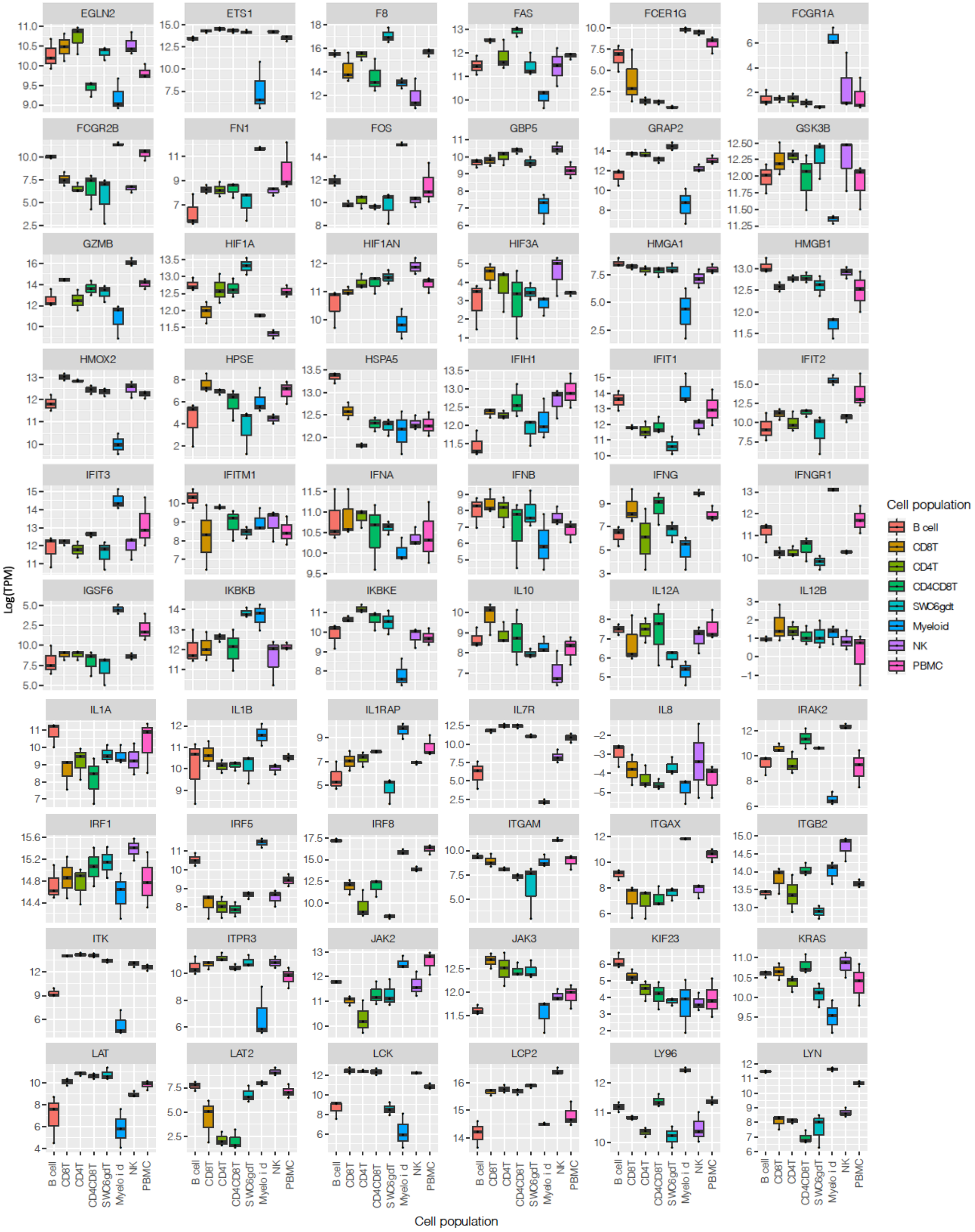
EGLN2
ETS1
F8
FAS
FCER1G
FCGR1A
FCGR2B
FN1
FOS
GBP5
GRAP2
GSK3B
GZMB
HIF1A
HIF1AN
HIF3A
HMGA1
HMGB1
HMOX2
HPSE
HSPA5
IFIH1
IFIT1
IFIT2
IFIT3
IFITM1
IFNA
IFNB
IFNG
IFNGR1
IGSF6
IKBKB
IKBKE
IL10
IL12A
IL12B
IL1A
IL1B
IL1RAP
IL7R
IL8
IRAK2
IRF1
IRF5
IRF8
ITGAM
ITGAX
ITGB2
ITK
ITPR3
JAK2
JAK3
KIF23
KRAS
LAT
LAT2
LCK
LCP2
LY96
LYN
Log(TPM)
Cell population
B cell
CD8T
CD4T
CD4CD8T
SWC6gdT
Myeloid
NK
PBMC
Cell population
B cell
CD8T
CD4T
CD4CD8T
SWC6gdt
Myeloid
NK
PBMC

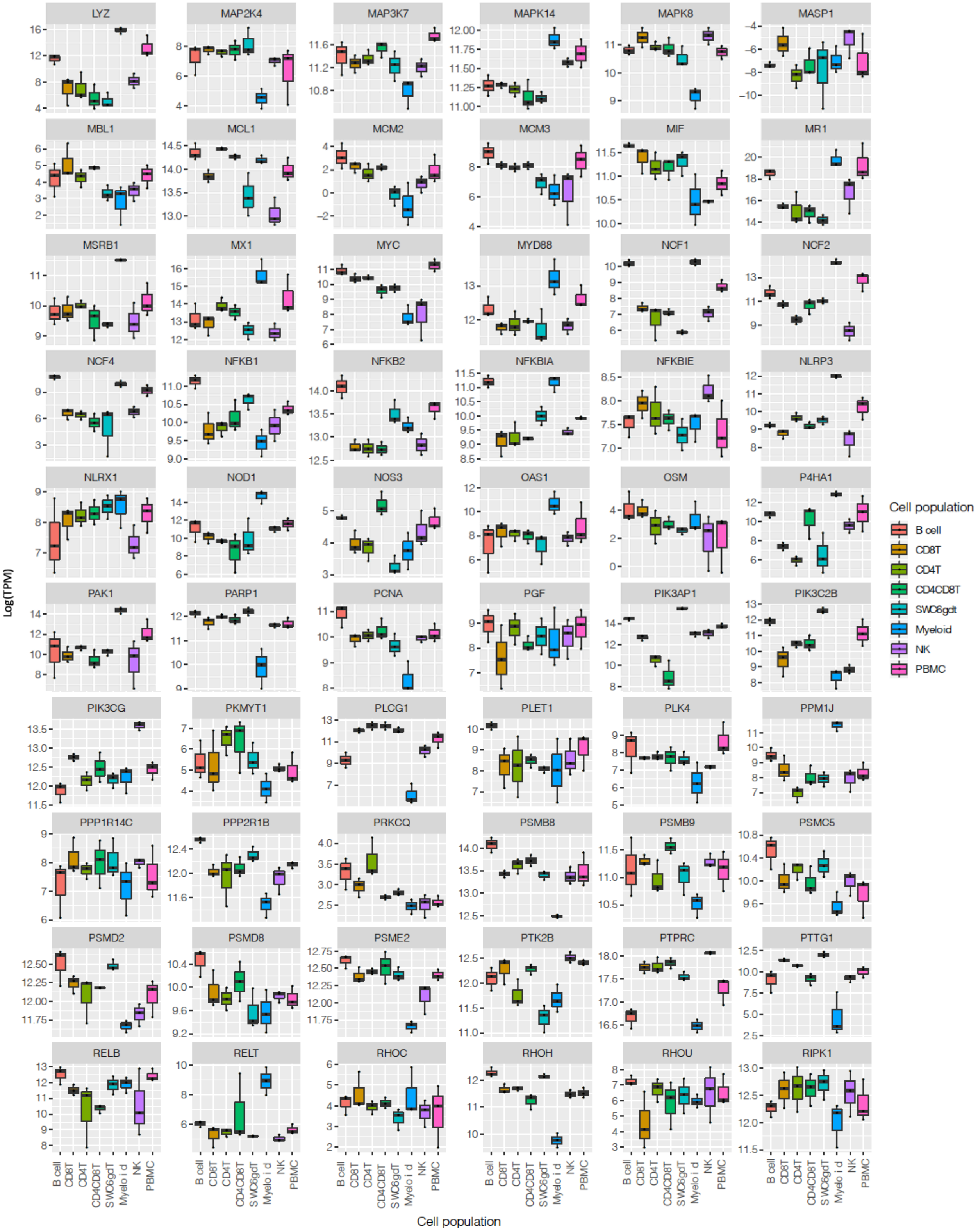
LYZ
MAP2K4
MAP3K7
MAPK14
MAPK8
MASP1
MBL1
MCL1
MCM2
MCM3
MIF
MR1
MSRB1
MX1
MYC
MYD88
NCF1
NCF2
NCF4
NFKB1
NFKB2
NFKBIA
NFKBIE
NLRP3
NLRX1
NOD1
NOS3
OAS1
OSM
P4HA1
PAK1
PARP1
PCNA
PGF
PIK3AP1
PIK3C2B
PIK3CG
PKMYT1
PLCG1
PLET1
PLK4
PPM1J
PPP1R14C
PPP2R1B
PRKCQ
PSMB8
PSMB9
PSMC5
PSMD2
PSMD8
PSME2
PTK2B
PTPRC
PTTG1
RELB
RELT
RHOC
RHOH
RHOU
RIPK1
Log(TPM)
Cell population
B cell
CD8T
CD4T
CD4CD8T
SWC6gdT
Myeloid
NK
PBMC
Cell population
B cell
CD8T
CD4T
CD4CD8T
SWC6gdt
Myeloid
NK
PBMC

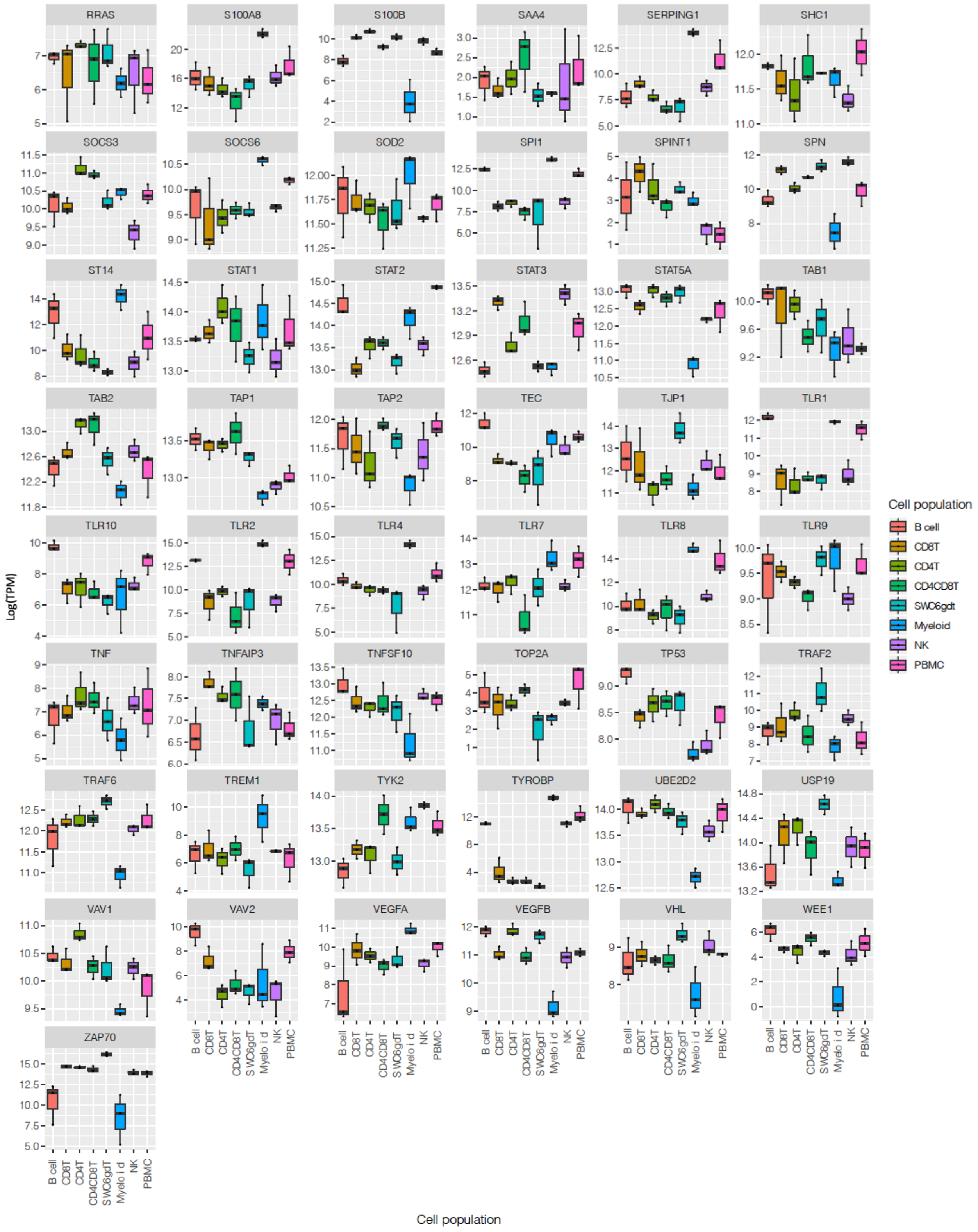


**Supplementary Figure 7.** RNA abundance of Nanostring selected genes. Box plot showing log(TPM) expression values of Nanostring selected genes of porcine sorted cells.

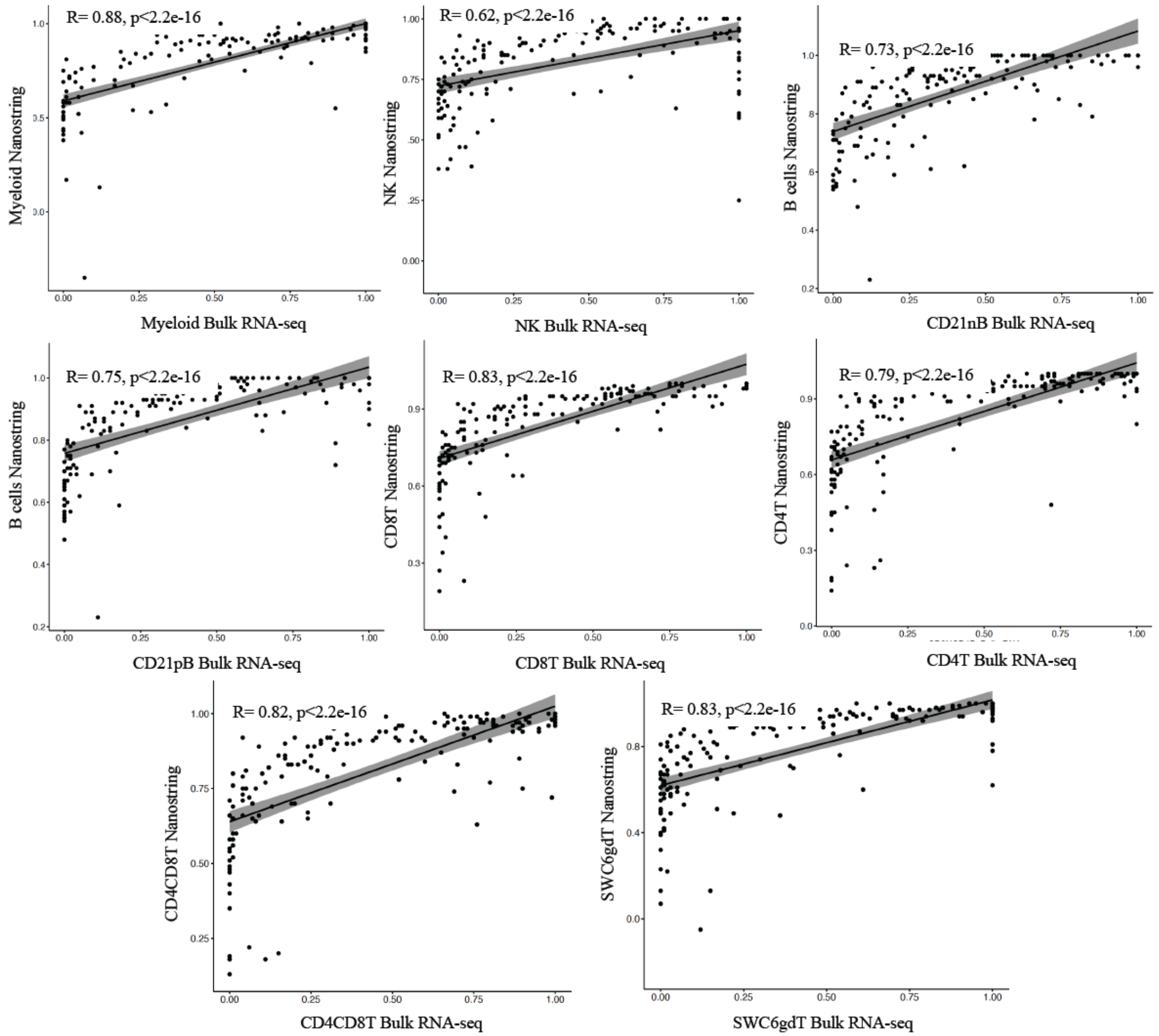


**Supplementary Figure 8.** Bulk RNA and Nanostring correlation. Spearman rank correlation of absolute normalize TPM values of common genes between bulk RNA-seq and Nanostring analysis.

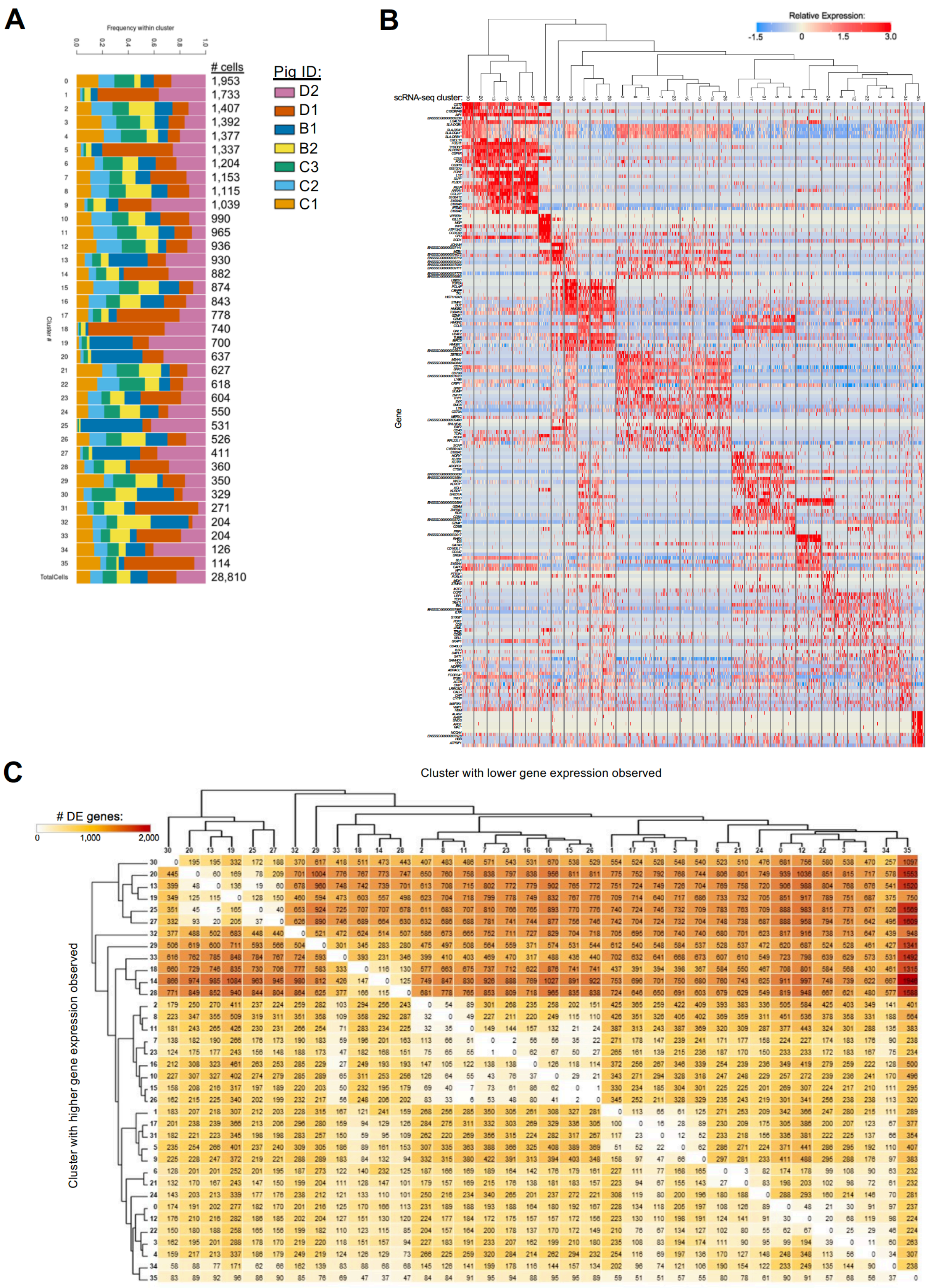
A
Frequency within cluster
0.0
0.2
0.4
0.6
0.8
1.0
### cells
Cluster #
0 1,953
1 1,733
2 1,407
3 1,392
4 1,377
5 1,337
6 1,204
7 1,153
8 1,115
9 1,039
10 990
11 965
12 936
13 930
14 882
15 874
16 843
17 778
18 740
19 700
20 637
21 627
22 618
23 604
24 550
25 531
26 526
27 411
28 360
29 350
30 329
31 271
32 204
33 204
34 126
35 114
TotalCells 28,810
Pig ID:
D2
D1
B1
B2
C3
C2
C1
B
Relative Expression:
-1.5
0
1.5
3.0
scRNA-seq cluster:
Gene
C
Cluster with lower gene expression observed
### DE genes:
0
1,000
2,000
Cluster with higher gene expression observed

**Supplementary Figure 9.** Unique characteristics of scRNA-seq clusters. A) Stacked barplot showing proportion of cells within each cluster derived from different PBMC samples. Bar color corresponds to the animal ID of a sample. Cluster IDs are shown on the left, while the number of cells within each cluster are listed on the right. Proportions of total cells are also listed in the bottom bar. **B)** Heatmap of the top DEGs within scRNA-seq clusters. Clusters are shown on the x-axis, along with a tree showing hierarchical clustering. Genes are listed on the y-axis. Up to the top five differentially expressed genes with highest positive logFC values for each cluster are listed. **C)** Heatmap showing the number of DEGs detected between scRNA-seq clusters by pairwise DGE analyses. Clusters are ordered on the x and y axis according to hierarchical relationships that were also shown in B. The cluster with higher gene expression for a pairwise comparison is shown on the y-axis, while the cluster with lower gene expression by pairwise comparison is shown on the x-axis. Seven PBMC samples used for scRNA-seq analysis were derived from each of three separate experiments (experiment B, n = 2; experiment C, n = 3; experiment D, n = 2) * Refer to 'Gene name replacement' methods.

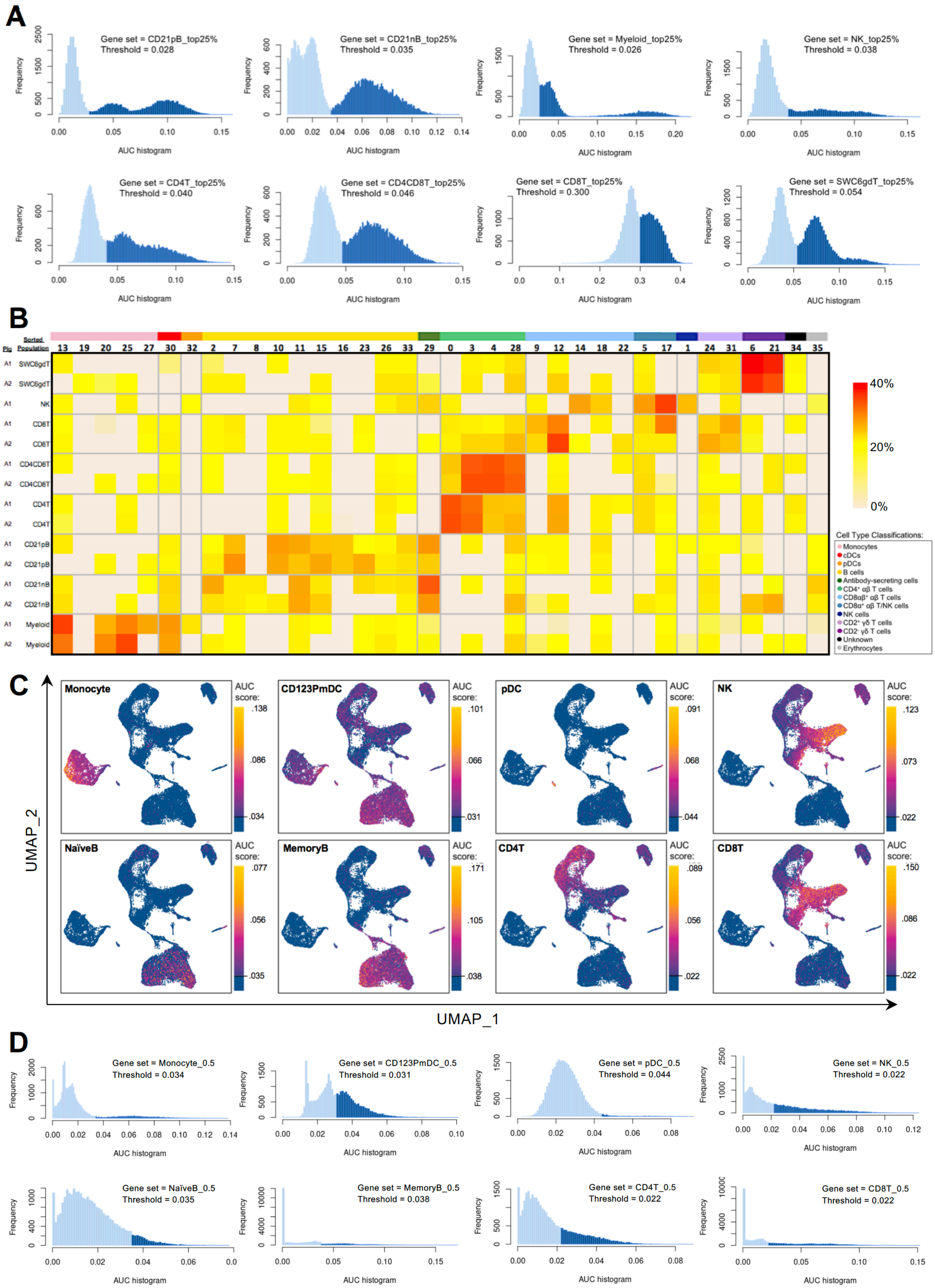
A
Gene set = CD21pB_top25%
Threshold = 0.028
Gene set = CD21nB_top25%
Threshold = 0.035
Gene set = Myeloid_top25%
Threshold = 0.026
Gene set = NK_top25%
Threshold = 0.038
Gene set = CD4T_top25%
Threshold = 0.040
Gene set = CD4CD8T_top25%
Threshold = 0.046
Gene set = CD8T_top25%
Threshold = 0.300
Gene set = SWC6gdT_top25%
Threshold = 0.054
Frequency
AUC histogram
B
Pig
Sorted Population
13 19 20 25 27 30 32 2 7 8 10 11 15 16 23 26 33 29 0 3 4 28 9 12 14 18 22 5 17 1 24 31 6 21 34 35
A1 SWC6gdT
A2 SWC6gdT
A1 NK
A1 CD8T
A2 CD8T
A1 CD4CD8T
A2 CD4CD8T
A1 CD4T
A2 CD4T
A1 CD21pB
A2 CD21pB
A1 CD21nB
A2 CD21nB
A1 Myeloid
A2 Myeloid
40%
20%
0%
Cell Type Classifications:
Monocytes
cDCs
pDCs
B cells
Antibody-secreting cells
CD4⁺ αβ T cells
CD8αβ⁺ αβ T cells
CD8α⁺ αβ T/NK cells
NK cells
CD2⁺ γδ T cells
CD2⁻ γδ T cells
Unknown
Erythrocytes
C
Monocyte
AUC score: .138 .086 .034
CD123PmDC
AUC score: .101 .066 .031
pDC
AUC score: .091 .068 .044
NK
AUC score: .123 .073 .022
NaïveB
AUC score: .077 .056 .035
MemoryB
AUC score: .171 .105 .038
CD4T
AUC score: .089 .056 .022
CD8T
AUC score: .150 .086 .022
UMAP_2
UMAP_1
D
Gene set = Monocyte_0.5
Threshold = 0.034
Gene set = CD123PmDC_0.5
Threshold = 0.031
Gene set = pDC_0.5
Threshold = 0.044
Gene set = NK_0.5
Threshold = 0.022
Gene set = NaïveB_0.5
Threshold = 0.035
Gene set = MemoryB_0.5
Threshold = 0.038
Gene set = CD4T_0.5
Threshold = 0.022
Gene set = CD8T_0.5
Threshold = 0.022
Frequency
AUC histogram

**Supplementary Figure 10.** Additional comparison between bulk and scRNA-seq data. **A)** Histogram of distribution of AUC scores for top 25% gene sets from porcine bulk RNA-seq sorted cell populations depicted in Figure 5 A-B. Frequency (y-axis) corresponds to the number of cells from the porcine scRNA-seq dataset having an AUC score value specified on the x-axis. Light blue bars represent cells falling below a threshold of positive AUC score detection, while dark blue bars represent cells with AUC scores above the threshold of detection used for plots in Figure 5A. Gene set and threshold of detection are provided for each respective histogram. **B)** Heatmap of CIBERSORTx deconvolution results. Porcine sorted cell bulk RNA-seq populations for each animal are listed on the y-axis, and porcine scRNA-seq clusters are listed on the x-axis. A color bar above scRNA-seq cluster IDs on the x-axis indicates the cell type classification, as according to Figure 4D. Heatmap color represents the proportion of bulk RNA-seq cells that correspond to each of the scRNA-seq clusters, with totals across rows for each bulk RNA-seq sample equaling 100%. **C)** Gene set enrichment scores calculated by AUCell analysis of enriched gene sets created from genes with high expression scores > 0.5 for each respective sorted population of human cells overlaid onto cells of the porcine scRNA-seq dataset visualized in 2-dimensional UMAP plot. Each dot represents a single cell. The color of the dot corresponds to the AUC score for each respective cell. Higher AUC scores correspond to greater gene set enrichment. A threshold for AUC score detection was set and indicated by a horizontal line on the gradient fill scale for each plot. The gene set mDC_0.5 was not included due to containing only a single gene used to calculate enrichment scores. **D)** Histogram of distribution of AUC scores from genes with high expression scores > 0.5 for each respective sorted population of human cells depicted in Figure 5 D & Supplementary Figure 10C. Frequency (y-axis) corresponds to the number of cells from the porcine scRNA-seq dataset having an AUC score value specified on the x-axis. Light blue bars represent cells falling below a threshold of positive AUC score detection, while dark blue bars represent cells with AUC scores above the threshold of detection used for plots in Figure 5A. Gene set and threshold of detection are provided for each respective histogram.

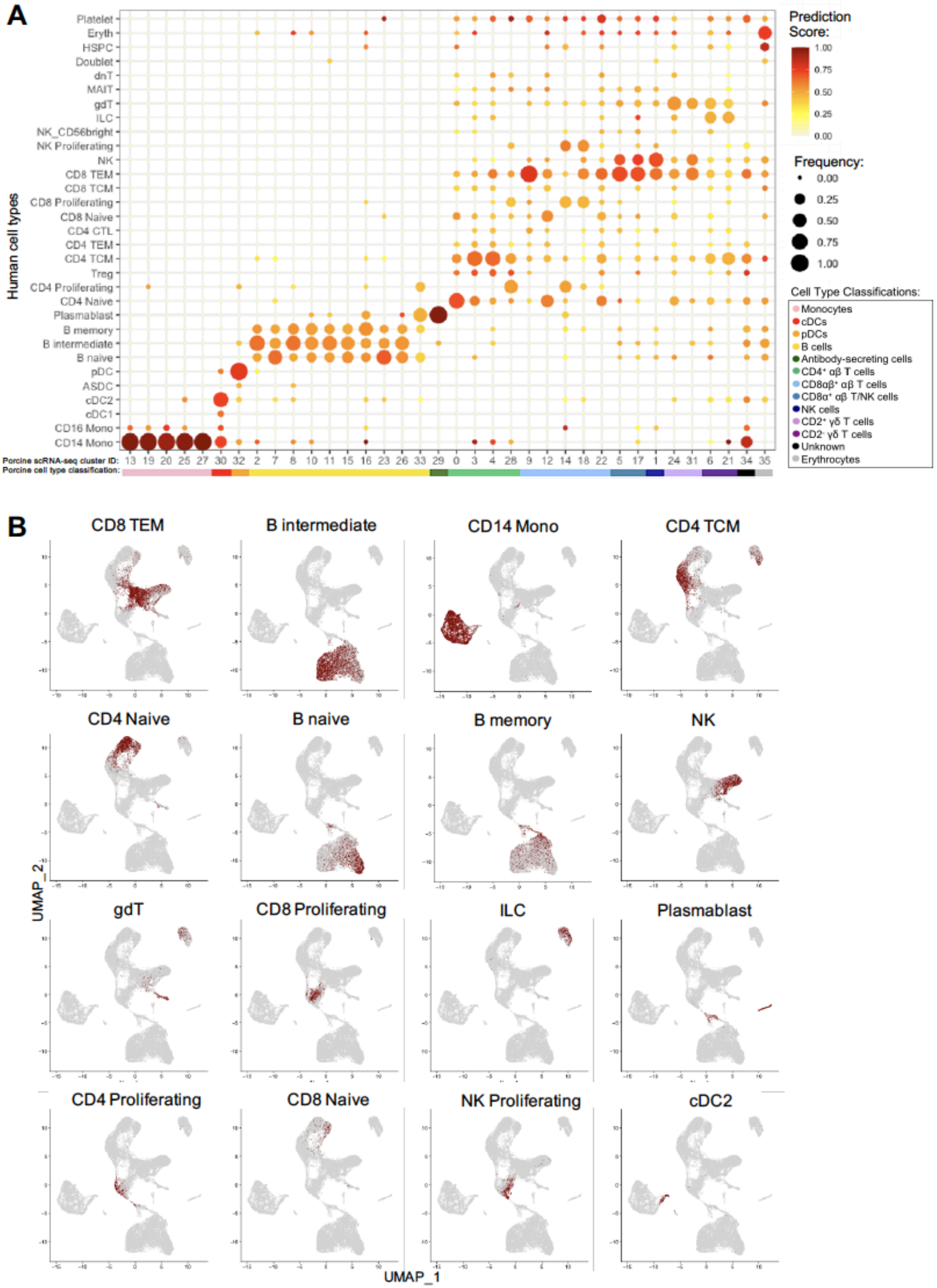
A
Prediction Score:
Frequency:
Human cell types
Platelet
Eryth
HSPC
Doublet
dnT
MAIT
gdT
ILC
NK_CD56bright
NK Proliferating
NK
CD8 TEM
CD8 TCM
CD8 Proliferating
CD8 Naive
CD4 CTL
CD4 TEM
CD4 TCM
Treg
CD4 Proliferating
CD4 Naive
Plasmablast
B memory
B intermediate
B naive
pDC
ASDC
cDC2
cDC1
CD16 Mono
CD14 Mono
Porcine scRNA-seq cluster ID:
Porcine cell type classification:
Cell Type Classifications:
Monocytes
cDCs
pDCs
B cells
Antibody-secreting cells
CD4⁺ αβ T cells
CD8αβ⁺ αβ T cells
CD8α⁺ αβ T/NK cells
NK cells
CD2⁺ γδ T cells
CD2⁻ γδ T cells
Unknown
Erythrocytes
B
CD8 TEM
B intermediate
CD14 Mono
CD4 TCM
CD4 Naive
B naive
B memory
NK
gdT
CD8 Proliferating
ILC
Plasmablast
CD4 Proliferating
CD8 Naive
NK Proliferating
cDC2
UMAP_2
UMAP_1

**Supplementary Figure 10.** Integration of porcine and human scRNA-seq datasets and transfer of human cell type labels. **A)** Prediction scores calculated to determine the confidence of cell type annotations. The human cell type specific frequency (size of the circle) and prediction score for that human cell type (color) are shown for each porcine scRNA-seq cluster. Porcine cell type classifications (color) are shown below the porcine scRNA-seq cluster IDs. **B)** The transferred human cell type labels are shown on the original porcine dataset UMAP. Each tile highlights a set of porcine cells annotated with their highest confidence level human cell type label in dark red.
